# Spatial sampling in human visual cortex is modulated by both spatial and feature-based attention

**DOI:** 10.1101/147223

**Authors:** D.M. van Es, J. Theeuwes, T. Knapen

## Abstract

Spatial attention changes the sampling of visual space. Behavioral studies suggest that feature-based attention modulates this resampling to optimize the attended feature’s sampling. We investigate this hypothesis by estimating spatial sampling in visual cortex while independently varying both feature-based and spatial attention.

Our results show that spatial and feature-based attention interacted: resampling of visual space depended on both the attended location and feature (color vs. temporal frequency). This interaction occurred similarly throughout visual cortex, regardless of an area’s overall feature preference. However, the interaction did depend on spatial sampling properties of voxels that prefer the attended feature. These findings are parsimoniously explained by variations in the precision of an attentional gain field.

Our results demonstrate that the deployment of spatial attention is tailored to the spatial sampling properties of units that are sensitive to the attended feature.

## Introduction

The resolution of the visual system is highest at the fovea and decreases gradually with increasing eccentricity But the visual system’s resolution is not fixed. Attention can be directed to a location in space and/or a visual feature, which temporarily improves perceptual sensitivity (Posner et al., 1980; Rossi and Paradiso, 1995; Found and Müller, 1996; Carrasco and Yeshurun, 1998; Yeshurun and Carrasco, 1999; Kumada, 2001; Saenz et al., 2003; Wolfe et al., 2003; Theeuwes and van der Burg, 2007) at the cost of reduced sensitivity for non-attended locations and features (Kastner and Pinsk, 2004; Pestilli and Carrasco, 2005; Wegener et al., 2008).

Attending a location in space increases activity in units representing the attended location, as shown by both electrophysiological (Luck et al., 1997; Reynolds et al., 2000) and fMRI studies (Tootell et al., 1998; Silver et al., 2005; Datta and DeYoe, 2009). In addition, spatial receptive fields were shown to shift toward an attended location in macaque MT+ (Womelsdorf et al., 2006) and V4 (Connor et al., 1997). Using fMRI to measure population receptive fields (pRFs; Dumoulin and Wandell, 2008; Dumoulin and Knapen, 2018), it was found that pRF shifts induced by spatial attention occur throughout human visual cortex (Klein et al., 2014; Kay et al., 2015; Sheremata and Silver, 2015; Vo et al., 2017), a process thought to improve visual resolution at the attended location (Anton-Erxleben and Carrasco, 2013; Kay et al., 2015; Vo et al., 2017). Such *spatial resampling* is understood as the result of an interaction between bottom-up sensory signals and a top-down attentional gain field (Womelsdorf et al., 2008; Klein et al., 2014; Miconi and VanRullen, 2016).

Feature-based attention, for example directed toward color or motion, selectively increases activity in those units that represent the attended feature, as evidenced by electrophysiological (Treue and Maunsell, 1996; Treue and Trujillo, 1999; McAdams and Maunsell, 2000; Maunsell and Treue, 2006; Müller et al., 2006; Zhang and Luck, 2009; Zhou and Desimone, 2011), fMRI (Saenz et al., 2002; Serences and Boynton, 2007; Jehee et al., 2011), and behavioral reports (Saenz et al., 2003; White and Carrasco, 2011). These studies consistently show that feature-based attention modulates processing irrespective of the attended stimulus’s spatial location. In addition, feature-based attention also appears to shift featural tuning curves toward the attended value, as reported by both electrophysiological (Motter, 1994; David et al., 2008) and fMRI studies (Çukur et al., 2013).

The similarity in the effects of feature-based and spatial attention on affected neural units suggests a common neural mechanism for both sources of attention (Hayden and Gallant, 2005; Cohen and Maunsell, 2011). Yet spatial attention necessitates retino-topically precise feedback (Miconi and VanRullen, 2016), while feature-based attention operates throughout the visual field (Maunsell and Treue, 2006). Studies investigating whether one source of attention potentiates the other generally find that interactions are either nonexistent or very weak at the earliest stages of processing (David et al., 2008; Hayden and Gallant, 2009; Patzwahl and Treue, 2009; Zhang and Luck, 2009), but emerge at later stages of visual processing (Hillyard and Munte, 1984; Handy et al., 2001; Bengson et al., 2012; Ibos and Freedman, 2016), and ultimately influence behavior (Kingstone, 1992; Kravitz and Behrmann, 2011; Leonard et al., 2015; White et al., 2015; Nordfang et al., 2017). In addition, the effects of feature-based compared to spatial attention arise earlier in time (Hopf et al., 2004; Hayden and Gallant, 2005; Andersen et al., 2011). This supports the idea that feature-based attention can direct spatial attention toward or away from specific locations containing attended or unattended features (Cohen and Shoup, 1997; Cepeda et al., 1998; Burnett et al., 2016). Especially when attention is endogenously cued, feature-based attention has been argued to influence the spatial resampling induced by spatial attention in order to optimize sampling of visual features for behavior (Yeshurun and Carrasco, 1998, 2000; Yeshurun et al., 2008; Barbot and Carrasco, 2017). Together, these studies suggest that feature-based attention can influence the spatial resampling resulting from spatial attention.

To investigate this hypothesis directly, we measured pRFs under conditions of differential spatial attention (i.e. toward fixation or the mapping stimulus) and feature-based attention (i.e. toward the mapping stimulus’s motion or color content). We first characterize how spatial attention influences the sampling of visual space, and subsequently investigate how feature-based attention modulates this spatial resampling. An explicit gain-field interaction model allowed us to formally capture the pRF position changes resulting from our attentional manipulations (Klein et al., 2014).

In brief, our results show that pRF changes are stronger when attending the stimulus’s color compared to temporal frequency content. These modulations occurred similarly thoughout the visual system, regardless of an area’s bottom-up feature preference (such as areas MT+ for temporal frequency and areas VO and hV4 for color (Liu and Wandell, 2005; Brouwer and Heeger, 2009, 2013)). We suggest that the larger degree of spatial resampling when attending color is related to finer spatial sampling in relatively color preferring voxels. In addition, we show that these feature-based attentional modulations can be explained by changes in the precision of the attentional gain field.

## Results

We first characterize the pattern of pRF parameter changes that resulted from the differential allocation of spatial attention (i.e. either toward fixation or the moving bar stimulus). We explain this pattern of results from an attentional gain field perspective. Then, we investigate how feature-based attention (i.e. either toward color or temporal frequency changes in the bar) modulated the pRF changes, and how this modulation relates to bottom-up feature preference.

### Region of interest definition

Figure 1A shows voxels’ *Attend Fixation* location preferences, by depicting color-coded polar angle coordinates on an inflated cortical surface for one example participant’s right hemisphere. We examined the relation between pRF eccentricity and size within each of the retinotopic regions, and performed further analyses on those regions that showed clear progressions of polar angle on the surface as well as positive size-eccentricity relations in all participants, as shown in Figure 1B (and Supplementary Figure 1). In addition, we created a *combined* ROI that pooled voxels across selected ROIs in order to evaluate pRF changes across the visual system.

**Figure 1:**
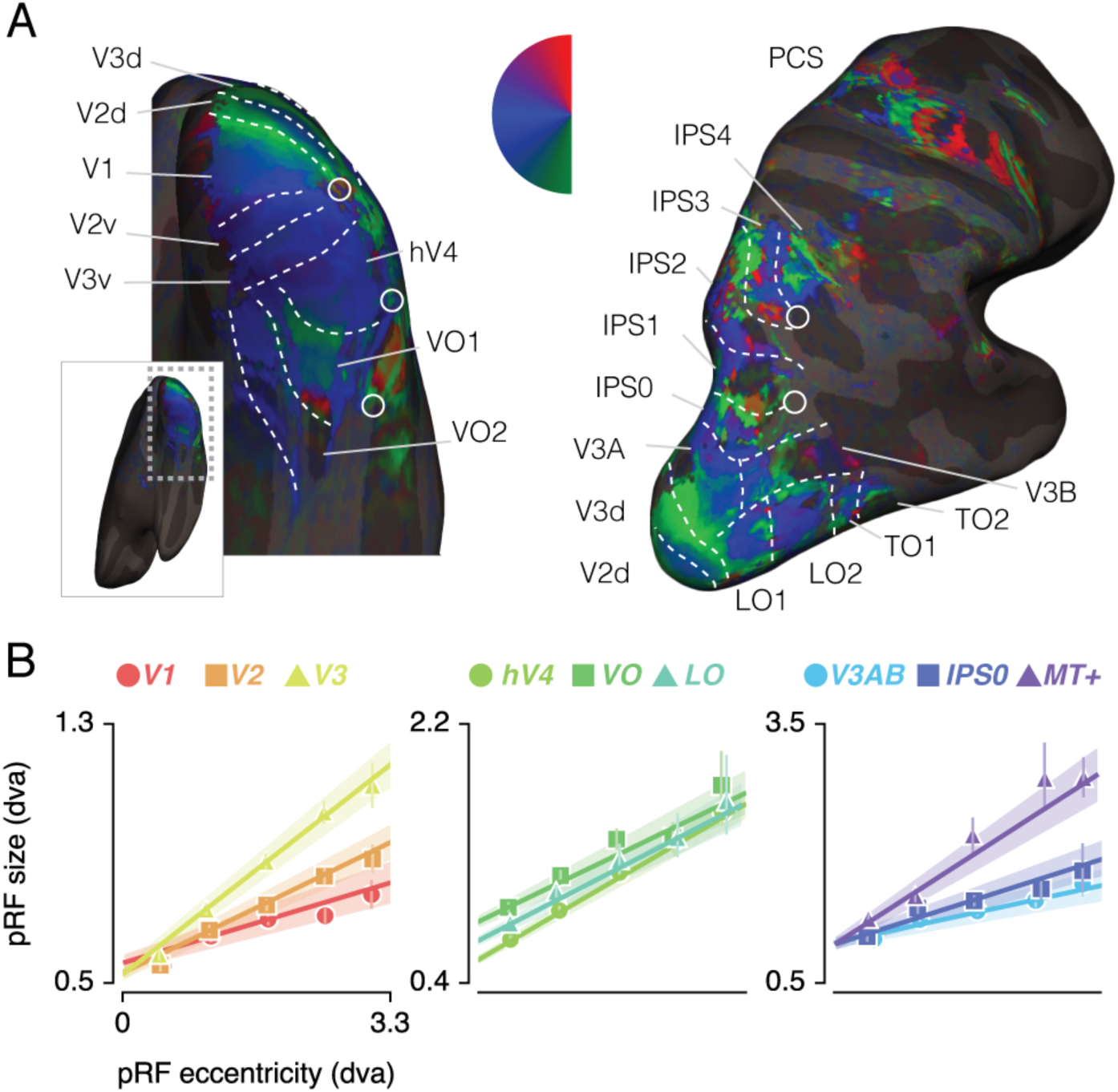
ROI definition A. *Attend Fixation* pRF polar angle maps for an example participant with retinotopic areas up to the intra-parietal sulcus defined by hand. B. *Attend Fixation* pRF size as a function of eccentricity for all areas that showed robust relationships across all participants. All error bars and shaded error regions denote 95% CI of data and linear fits respectively across voxels.

### pRF changes induced by spatial attention

In order to quantify pRF changes resulting from differential allocation of spatial attention we created an *Attend Stimulus* condition by averaging pRF parameters between the *Attend Color* and *Attend Temporal Frequency* (henceforth: *Attend TF*) conditions. To inspect how spatial attention affected pRF positions, we plotted a vector from the *Attend Fixation* to the *Attend Stimulus* pRF position (Figure 2A and B and Supplementary Figure 2). Visual inspection of these pRF position shifts shows both increasing shift magnitude up the visual hierarchy and shifts occuring mainly along the radial dimension (i.e. toward or away from the fovea).

**Figure 2:**
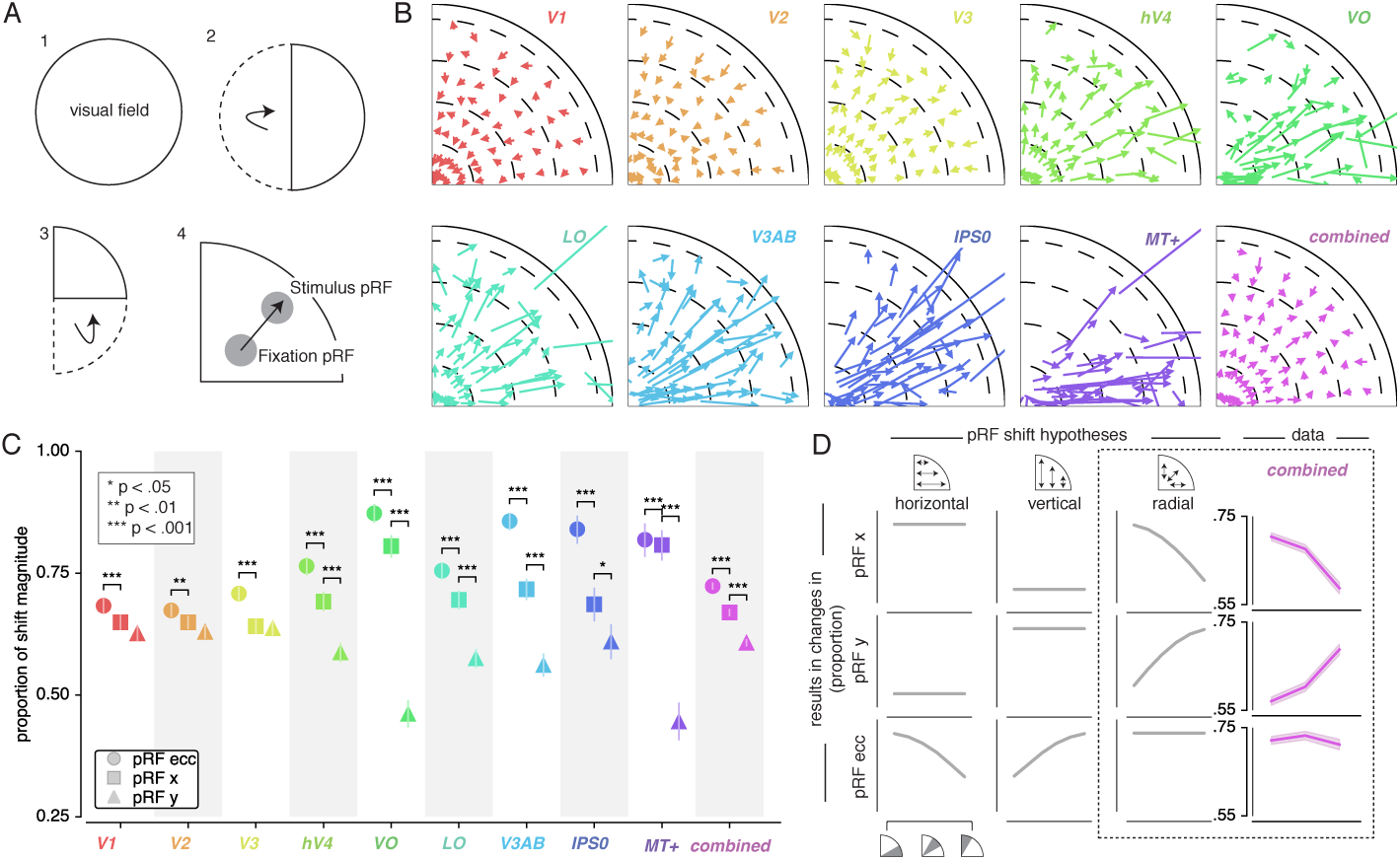
Effect of attention on pRF position. A. Plotting strategy. For pRF shift visualizations, all pRF positions are mirrored into one quadrant of the visual field. Then, vectors representing the shift of pRF centers between conditions were drawn from the *Attend Fixation* to the *Attend Stimulus* pRF position. B. Shift vectors as described in A. pRF shift magnitude increased up the visual hierarchy, and shifts appear to occur mainly in the radial direction (i.e. changes in pRF eccentricity). Dotted lines demarcate eccentricity bins used in subsequent analyses. C. Changes in pRF position in the horizontal, vertical, and radial directions as a proportion of the length of the shift vectors, as depicted in B. The magnitude of pRF shifts is consistently best described by changes in pRF eccentricity. D. pRF x, y and eccentricity position shifts plotted as a function of polar angle, for different shift direction hypotheses. The data closely matches the radial shift direction hypothesis, showing strongest pRF x shifts close to the horizontal meridian, strongest pRF y shifts close to the vertical meridian and strong pRF eccentricity changes across all polar angles. In C single, double, and triple asterisks indicate significant differences with FDR corrected p < .05, < .01 and < .001 respectively.

**Table 1:** Statistics corresponding to Figure 2C on pRF shift direction ratios. P-values reflect proportion of bootstrapped differences that are different from 0. Single, double and triple asterisks indicate FDR corrected significance of <.05, <.01 and <.001 respectively. FDR correction performed over all p-values in this table simultaneously.

| x>y |  |  |  |  | ecc>x |  |  |  |
| --- | --- | --- | --- | --- | --- | --- | --- | --- |
| ROI | N | p | $\Delta$ ratio | cohen's d | N | p | $\Delta$ ratio | cohen's d |
| V1 | 2176 | .098 | .02 | .04 | 2176 | <.001*** | .03 | .09 |
| V2 | 2752 | .102 | .02 | .03 | 2752 | .002** | .02 | .06 |
| V3 | 2320 | .773 | .00 | .01 | 2318 | <.001*** | .07 | .18 |
| hV4 | 1201 | <.001*** | .10 | .18 | 1201 | <.001*** | .07 | .20 |
| VO | 582 | <.001*** | .34 | .67 | 573 | <.001*** | .08 | .37 |
| LO | 1397 | <.001*** | .12 | .20 | 1394 | <.001*** | .06 | .16 |
| V3AB | 880 | <.001*** | .16 | .27 | 873 | <.001*** | .15 | .46 |
| IPS0 | 313 | .023* | .08 | .14 | 313 | <.001*** | .16 | .61 |
| MT+ | 325 | <.001*** | .36 | .68 | 306 | <.001*** | .04 | .28 |
| combined | 11946 | <.001*** | .06 | .10 | 11940 | <.001*** | .05 | .15 |

**Table 2:**
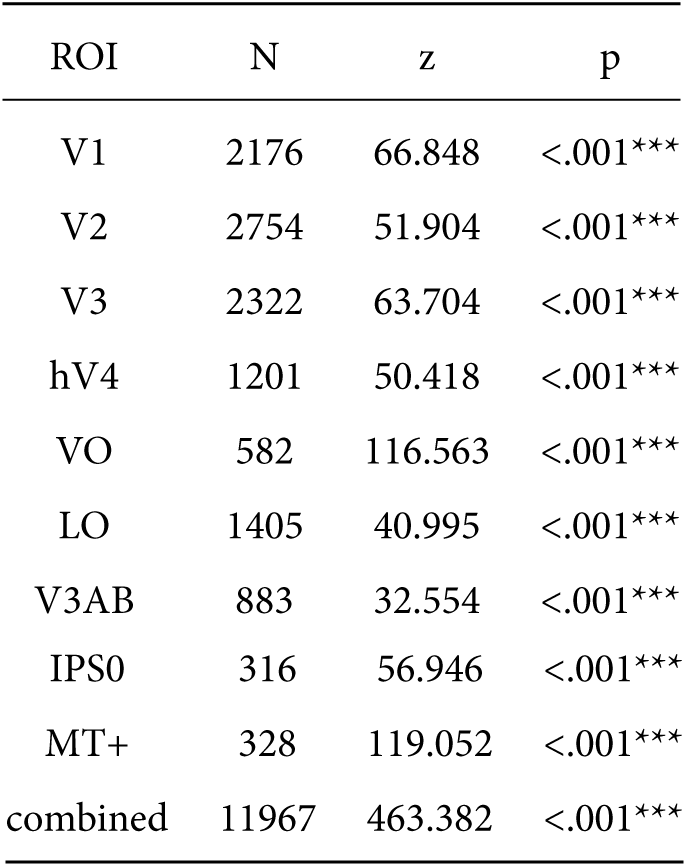
Statistics corresponding to Figure 2 on uniformity of polar angle distributions. P-values test whether pRFs are distributed non-uniformly over polar angle (Rayleigh test). Triple asterisks indicate FDR corrected significance of <.001. FDR correction performed over all p-values in this table simultaneously.

| ROI | N | z | p |
| --- | --- | --- | --- |
| V1 | 2176 | 66.848 | <.001*** |
| V2 | 2754 | 51.904 | <.001*** |
| V3 | 2322 | 63.704 | <.001*** |
| hV4 | 1201 | 50.418 | <.001*** |
| VO | 582 | 116.563 | <.001*** |
| LO | 1405 | 40.995 | <.001*** |
| V3AB | 883 | 32.554 | <.001*** |
| IPS0 | 316 | 56.946 | <.001*** |
| MT+ | 328 | 119.052 | <.001*** |
| combined | 11967 | 463.382 | <.001*** |

#### pRF shift direction

This latter observation seems at apparent odds with a recent study reporting pRFs to mainly shift in the horizontal direction (Sheremata and Silver, 2015). To quantify the observed direction of pRF shifts we computed the ratio of shifts in the radial, horizontal, and vertical directions (see Figure 2C and Supplementary Figure 3). In line with the data of Sheremata and Silver (2015), we find that changes of pRF horizontal location consistently better describe the overall shifts than do changes of pRF vertical location in all ROIs except V1/2/3 (p’s < .05, see Supplementary Tables 1 and 10). We also find that pRF shifts are described even better by shifts in the radial dimension (i.e. changes in eccentricity) compared to shifts in the horizontal direction in all ROIs (p’s < .01, see Supplementary Tables 1 and 11). Figure 2D is intended to ease interpretation of these results. It depicts how different hypotheses regarding the underlying directionality of pRF shifts (i.e. horizontal, vertical, or radial - i.e. foveopetal/foveofugal) translate into changes in measured pRF x, y, and eccentricity as a function of quarter visual field polar angle (i.e. from vertical to horizontal meridian). For example, if pRFs shift primarily in the radial direction (right hypothesis column, Figure 2D), this would result in strongest pRF x-direction changes close to the horizontal meridian and strongest pRF y-direction changes close to the vertical meridian. pRF eccentricity changes however, would show no dependence on polar angle. Figure 2D, right column, shows that the data (*combined* ROI) corresponds most to the radial shift hypothesis. To quantify this visual intuition, we compared the slopes of the change in pRF *x* and *y* over polar angle by binning polar angle into three bins and comparing the first and last bins (i.e. horizontal and vertical meridian respectively). This showed that, compared to the slope of pRF *y* change over polar angle, the slope of pRF *x* change was more negative (p < .001, cohen’s d = 0.677, N = 11946, see Supplementary Figure 4). This pattern of results can only be explained by pRFs shifting in the radial direction. Visual field coverage is known to be non-uniform such that the horizontal meridian is overrepresented at both subcortical (Schneider et al., 2004) and cortical (Swisher et al., 2007; Silva et al., 2017) levels and was also clearly present in our data (Rayleigh tests for non-uniformity in ROIs separately, p’s < .001, see Supplementary Tables 2, 12 and 13). This means that shifts that occur exclusively in the radial dimension appear as a dominance of horizontal compared to vertical shifts when averaging over the visual field.

#### pRF changes across eccentricity

To further inspect the attention-induced radial shifts described above, we plotted the difference between *Attend Stimulus* and *Attend Fixation* pRF eccentricity for each of four *Attend Fixation* pRF eccentricity bins (Figure 3A and Supplementary Figure 5). The *combined* ROI shows that overall, parafoveal pRFs shifted away from the fovea, while peripheral pRFs shifted toward the fovea. These outward shifting parafoveal pRFs are found in all other ROIs except V1 and V2, whereas the inward shifting peripheral pRFs are also present in V1, V2 and V3 (see Supplementary Tables 3, 4, and 14-17).

**Figure 3:**
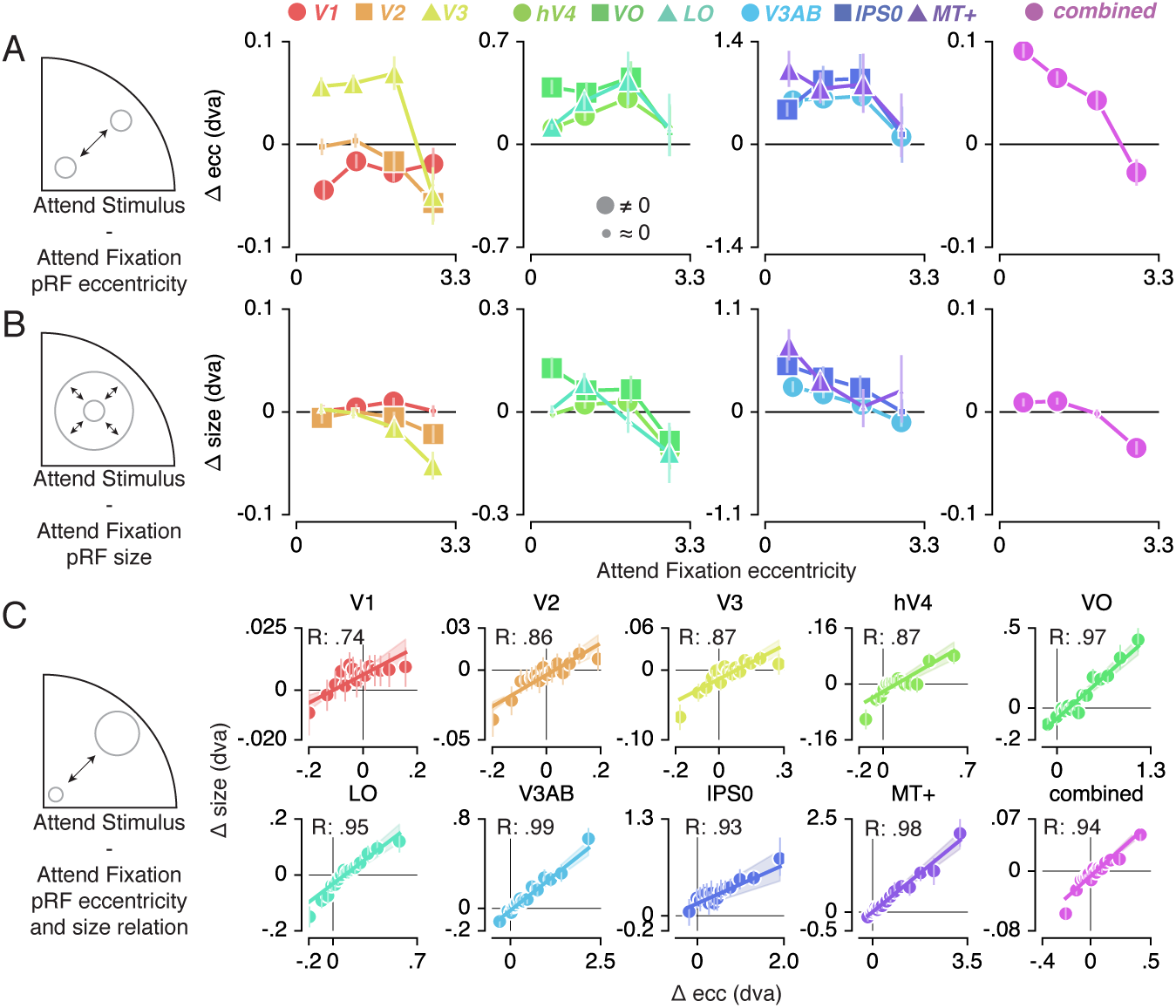
Effect of attention on pRF eccentricity and size. Difference between *Attend Stimulus* and *Attend Fixation* pRF eccentricity (A) and size (B) as a function of *Attend Fixation* eccentricity. Overall, parafoveal pRFs tend to shift away from the fovea and increase in size, while peripheral pRFs tend to shift toward the fovea and decrease in size. C. Changes in pRF eccentricity and size were strongly correlated in all ROIs. In A and B, markers are increased in size when bootstrapped distributions differ from 0 with FDR corrected p < .05. In C markers’ errorbar denotes 95% CI of data over voxels and shaded error regions denote 95% CI of linear fit parameters over bins.

In addition to pRF position changes, we also inspected changes in pRF size induced by differences in spatial attention as a function of *Attend Fixation* pRF eccentricity (Figure 3B and Supplementary Figure 6). Overall, parafoveal pRFs increased in size, while peripheral pRFs decreased in size. These expanding parafoveal pRFs were present in all ROIs except V2/3, whereas shrinking peripheral pRFs were found in all ROIs except V1, MT+ and IPSO (see Supplementary Tables 5, 6, and 18-21). Overall, this pattern of results is strikingly similar to the changes in pRF eccentricity described above. In fact, the changes in pRF size and eccentricity were strongly correlated in all ROIs (Figure 3C; Supplementary Figure 6; Pearson R over 20 5-percentile bins between .74 and .99, p’s < .001, see Supplementary Tables 7 and 22). Together, these results show that attention to the stimulus caused parafoveal pRFs to shift away from the fovea and increase in size, whereas peripheral pRFs shifted toward the fovea and decreased in size.

#### Formal account for observed pattern of pRF shifts

In order to provide a mechanistic explanation for the complex pattern of pRF shifts described above, we modeled our results using a multiplicative Gaussian gain field model (Womelsdorf et al., 2008; Klein et al., 2014). We adapted this framework to work in conditions where attention shifted over space as a function of time (see Methods). In brief, this modeling procedure used the *Attend Fixation* pRF, one attentional gain field at fixation and another convolved with the stimulus in order to predict the *Attend Stimulus* pRF position. We determined optimal attentional gain field sizes by minimizing the difference between observed and predicted *Attend Stimulus* pRF positions in the quadrant visual field format of Figure 2B. Figure 4A (and Supplementary Figure 7) illustrates that model predictions closely followed the data, thereby accurately reproducing radially shifting pRFs. Examining the predicted change in pRF eccentricity as a function of eccentricity (i.e. the dominant pRF shift direction; Figure 4B and Supplementary Figure 8) showed that the model was able to capture widely varying eccentricity change profiles across ROIs using very similar attentional gain field sizes (Figure 4C). This shows that a common attentional influence can result in very different pRF shift patterns, which then necessarily depend on differential spatial sampling properties across ROIs (i.e. distribution of pRF sizes and positions). In sum, these results show that the attentional gain field model provides a parsimonious and powerful account for the variety of pRF shift patterns across ROIs.

**Figure 4:**
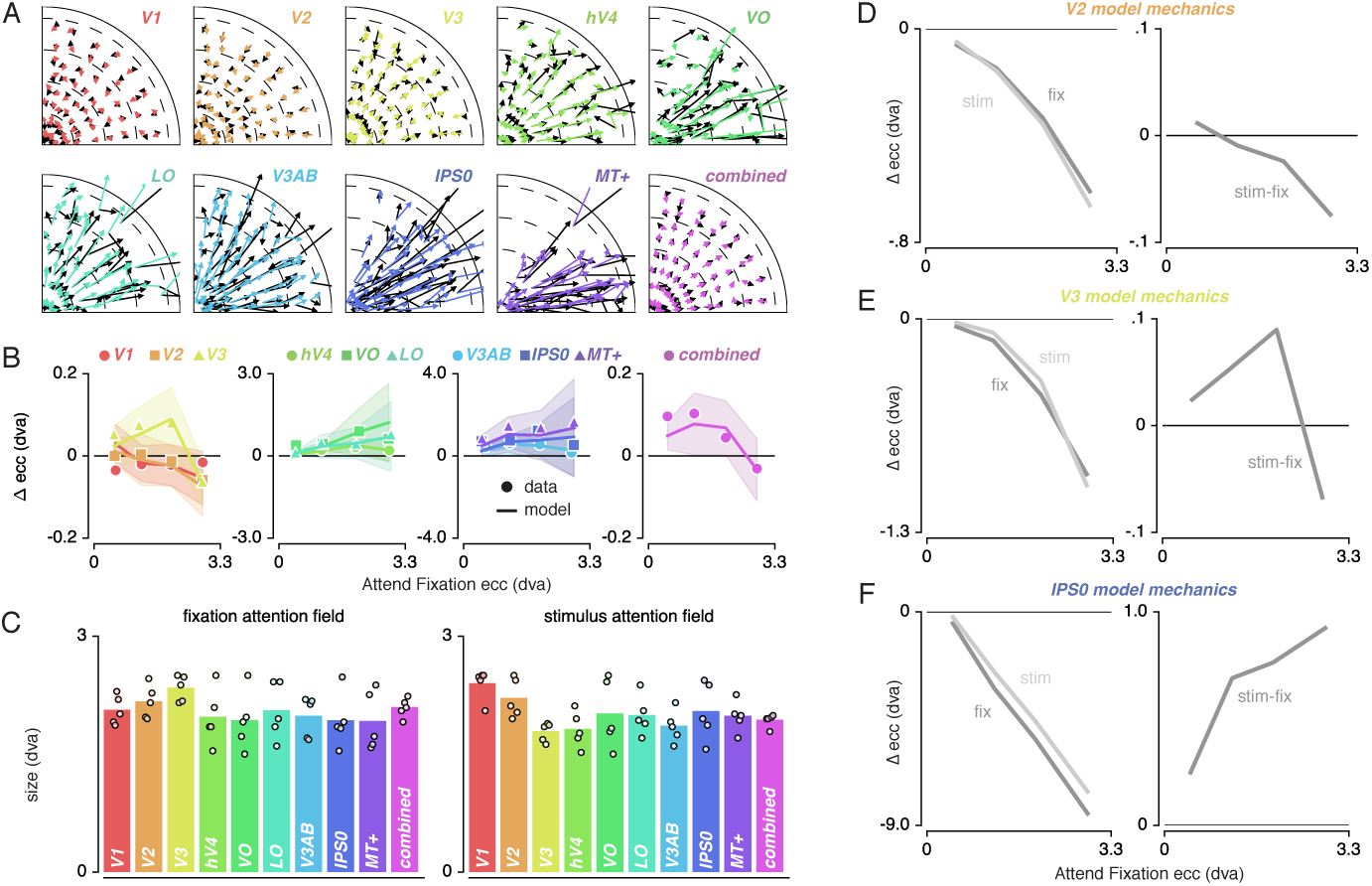
An attentional gain field model showed that the changes in pRF position can be described by a Gaussian interaction process. A. Observed (black) and predicted (color) pRF shifts. B. Observed and predicted changes in pRF eccentricity (the main pRF shift direction) as a function of eccentricity. Markers depict data and lines the corresponding attentional gain field model fit. C. Fitted fixation and stimulus attentional gain field sizes. Dots depict individual subjects, and the bar the average across subjects. D-F. Left panels depict changes in eccentricity induced by attending fixation (dark gray lines) and by attending the stimulus (light gray lines). Although both sources of attention cause a pull toward the fovea in all ROIs, relative shift magnitude differs across eccentricity. The right panels show how the difference between both spatial attention conditions results in the patterns as observed in B. In B, Error bars denote 95% CIs over subjects. Plotting conventions as in Figure 2.

We further investigated how the model was able to reproduce the eccentricity-dependent eccentricity changes we reported above. For this, we inspected pRF shifts induced by attending either fixation or the stimulus relative to the stimulus drive (i.e. the pRF outside the influence of attention derived from the model). For illustrative purposes, we here display results for V2, V3 and IPSO as these areas showed marked differences in their eccentricity change profile (Figure 4D-F). The left panels of each figure reveal the effects of attending fixation and the stimulus separately. This shows that both sources of spatial attention pull the measured pRFs toward the fovea, albeit with differing relative magnitudes across eccentricity. The right panel of each figure shows that the resulting difference between attending fixation and the stimulus constitutes the eccentricity dependent patterns observed in the data (Figure 4B).

Together, these analyses show that existing multiplicative gain field models of attention can be extended to predict pRF shifts in situations where spatial attention shifts over time. Additionally it confirms, extends, and quantifies earlier reports showing that the precision of the attentional gain field is similar across the visual hierarchy (Klein et al., 2014).

### Feature-based attentional modulation

Having established (1) the pattern of changes in spatial sampling (i.e. changes in pRF size and eccentricity) resulting from differential allocation of spatial attention and (2) a mechanistic explanation of these changes, we next examined how this pattern was modulated by differences in feature-based attention. Figure 5A (and Supplementary Figure 9) shows how pRF eccentricity and size are differentially affected by attending color or temporal frequency within the stimulus for the *combined* ROI. This illustrates that while both tasks caused similar pRF changes, these effects were generally more pronounced when attending color.

**Figure 5:**
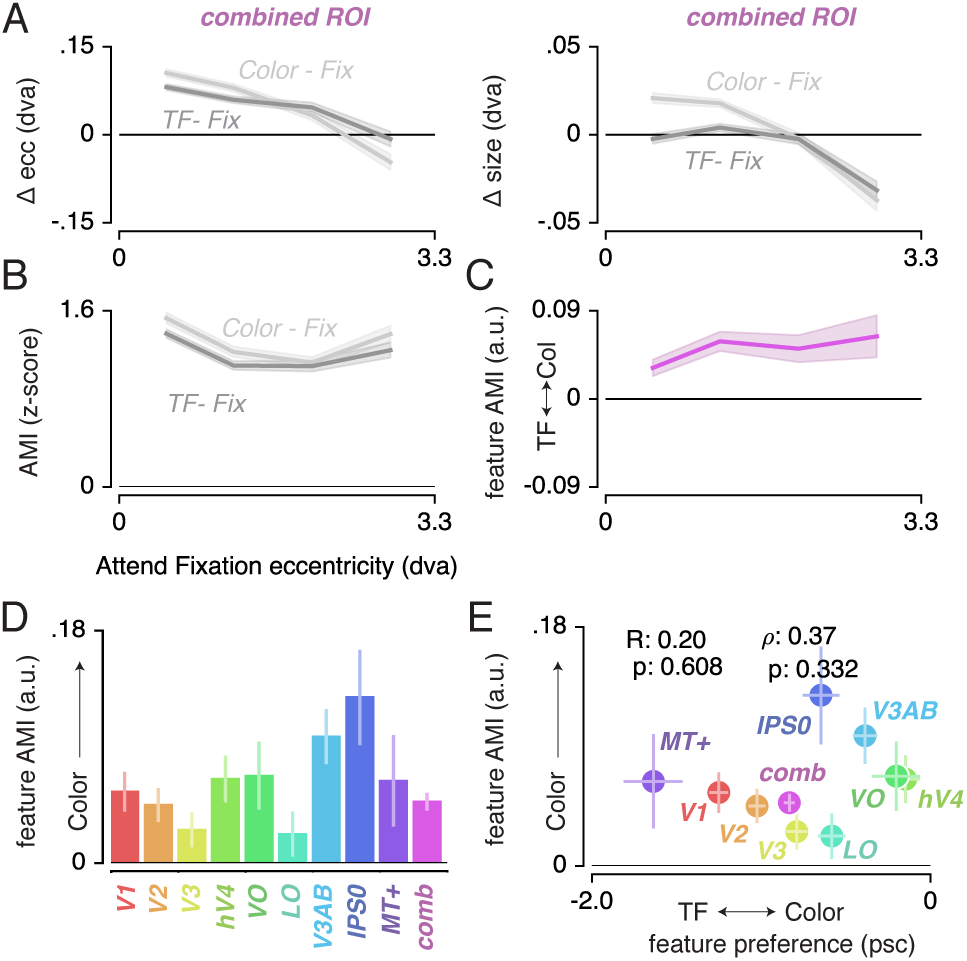
Feature-based attentional modulation across ROIs. A. Differences in pRF eccentricity and size relative to the *Attend Fixation* condition, for both the *Attend Color* and *Attend TF* condition. The changes in both eccentricity and size are more pronounced when attending changes in color versus TF changes in the bar. B. The Attentional Modulation Index (AMI) combines eccentricity and size changes to form one robust index of spatial attention and is greater when attending color compared to TF. C. The feature AMI quantifies this difference. Positive values of this feature AMI across eccentricity confirm stronger pRF modulations when attending color compared to TF. D. Average feature AMI for each ROI, extending greater observed pRF modulations when attending color compared to TF to all individual ROIs. E. Average feature AMI as a function of average feature preference across ROIs. Feature preference increases with higher color compared to TF preference. Although hV4 and VO are relatively sensitive to color and MT+ is relatively sensitive to TF, feature AMI is comparable in these areas. Errorbars denote 95% CI over voxels.

**Table 3:** Statistics corresponding to Figure 3A on pRF eccentricity changes. P-values reflect whether bootstrapped distribution is different from 0, for each ROI and each eccentricity bin (bin 3 and 4 in table below). Single, double and triple asterisks indicate FDR corrected significance of <.05, <.01 and <.001 respectively. FDR correction performed over all p-values in this <u>table simultaneously.</u>

| ecc bin | 1 |  |  |  | 2 |  |  |  |
| --- | --- | --- | --- | --- | --- | --- | --- | --- |
| ROI | N | p | $\Delta$ ecc | cohen's d | N | p | $\Delta$ ecc | cohen's d |
| V1 | 509 | <.001*** | -.044 | -.44 | 749 | <.001*** | -.016 | -.20 |
| V2 | 862 | 0.504 | -.002 | -.02 | 931 | .204 | .004 | .04 |
| V3 | 920 | <.001*** | .056 | .49 | 747 | <.001*** | .059 | .58 |
| hV4 | 693 | <.001*** | .109 | .67 | 359 | <.001*** | .194 | .64 |
| VO | 313 | <.001*** | .389 | 1.04 | 162 | <.001*** | .352 | .85 |
| LO | 1047 | <.001*** | .121 | .62 | 263 | <.001*** | .295 | .60 |
| V3AB | 220 | <.001*** | .600 | .86 | 348 | <.001*** | .623 | .87 |
| IPS0 | 170 | <.001*** | .473 | .91 | 86 | <.001*** | .876 | 1.05 |
| MT+ | 186 | <.001*** | 1.015 | .68 | 103 | <.001*** | .759 | .66 |
| combined | 4920 | <.001*** | .091 | .45 | 3748 | <.001*** | .065 | .37 |

**Table 4:**
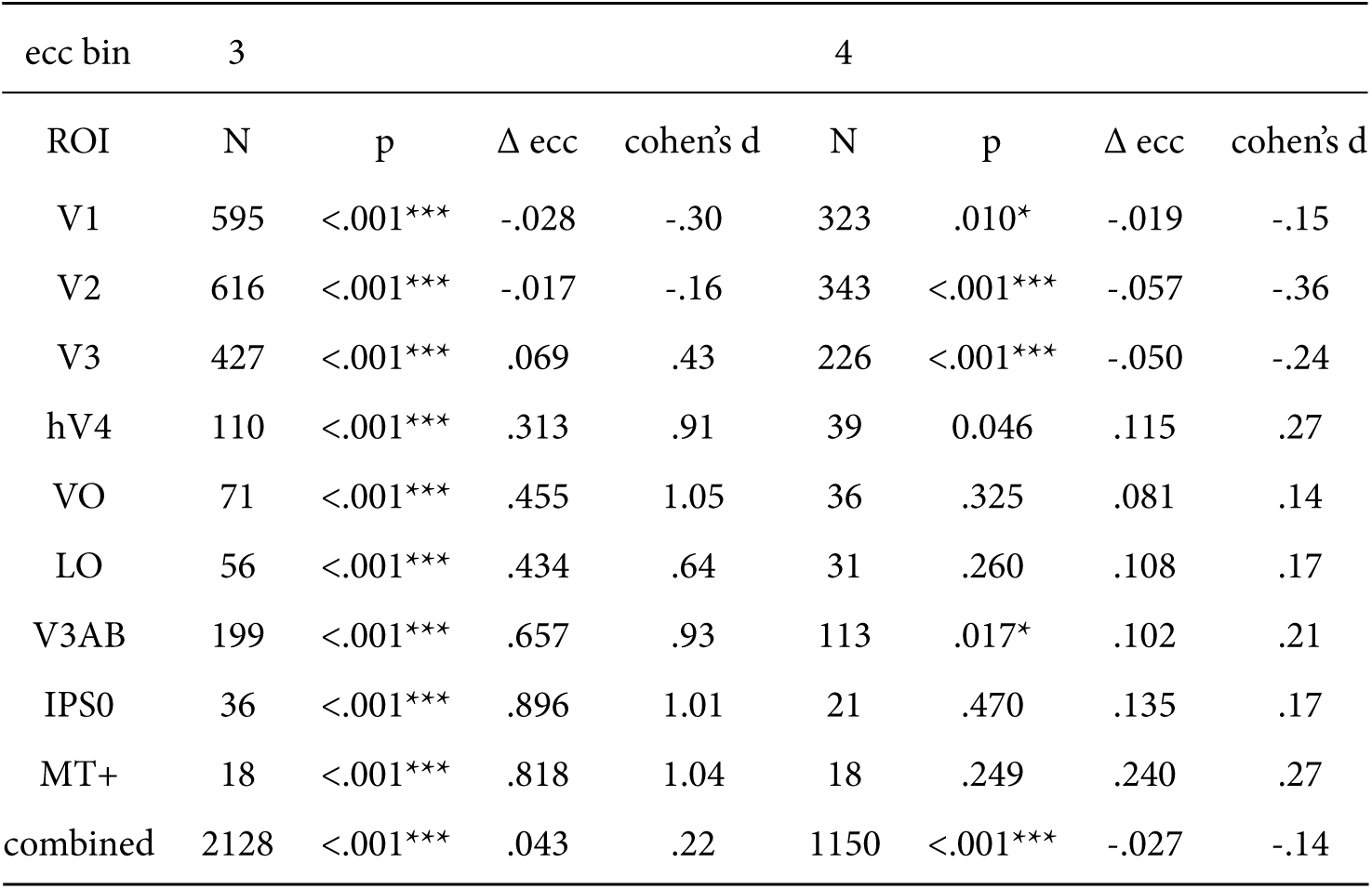
Statistics corresponding to Figure 3A on pRF eccentricity changes. P-values reflect whether bootstrapped distribution is different from 0, for each ROI and each eccentricity bin (bin 1 and 2 in table above). Single, double and triple asterisks indicate FDR corrected significance of <.05, <.01 and <.001 respectively. FDR correction performed overall p-values in this <u>table simultaneously.</u>

| ecc bin | 3 |  |  |  | 4 |  |  |  |
| --- | --- | --- | --- | --- | --- | --- | --- | --- |
| ROI | N | p | $\Delta$ ecc | cohen's d | N | p | $\Delta$ ecc | cohen's d |
| V1 | 595 | <.001*** | -.028 | -.30 | 323 | .010* | -.019 | -.15 |
| V2 | 616 | <.001*** | -.017 | -.16 | 343 | <.001*** | -.057 | -.36 |
| V3 | 427 | <.001*** | .069 | .43 | 226 | <.001*** | -.050 | -.24 |
| hV4 | 110 | <.001*** | .313 | .91 | 39 | 0.046 | .115 | .27 |
| VO | 71 | <.001*** | .455 | 1.05 | 36 | .325 | .081 | .14 |
| LO | 56 | <.001*** | .434 | .64 | 31 | .260 | .108 | .17 |
| V3AB | 199 | <.001*** | .657 | .93 | 113 | .017* | .102 | .21 |
| IPS0 | 36 | <.001*** | .896 | 1.01 | 21 | .470 | .135 | .17 |
| MT+ | 18 | <.001*** | .818 | 1.04 | 18 | .249 | .240 | .27 |
| combined | 2128 | <.001*** | .043 | .22 | 1150 | <.001*** | -.027 | -.14 |

**Table 5:**
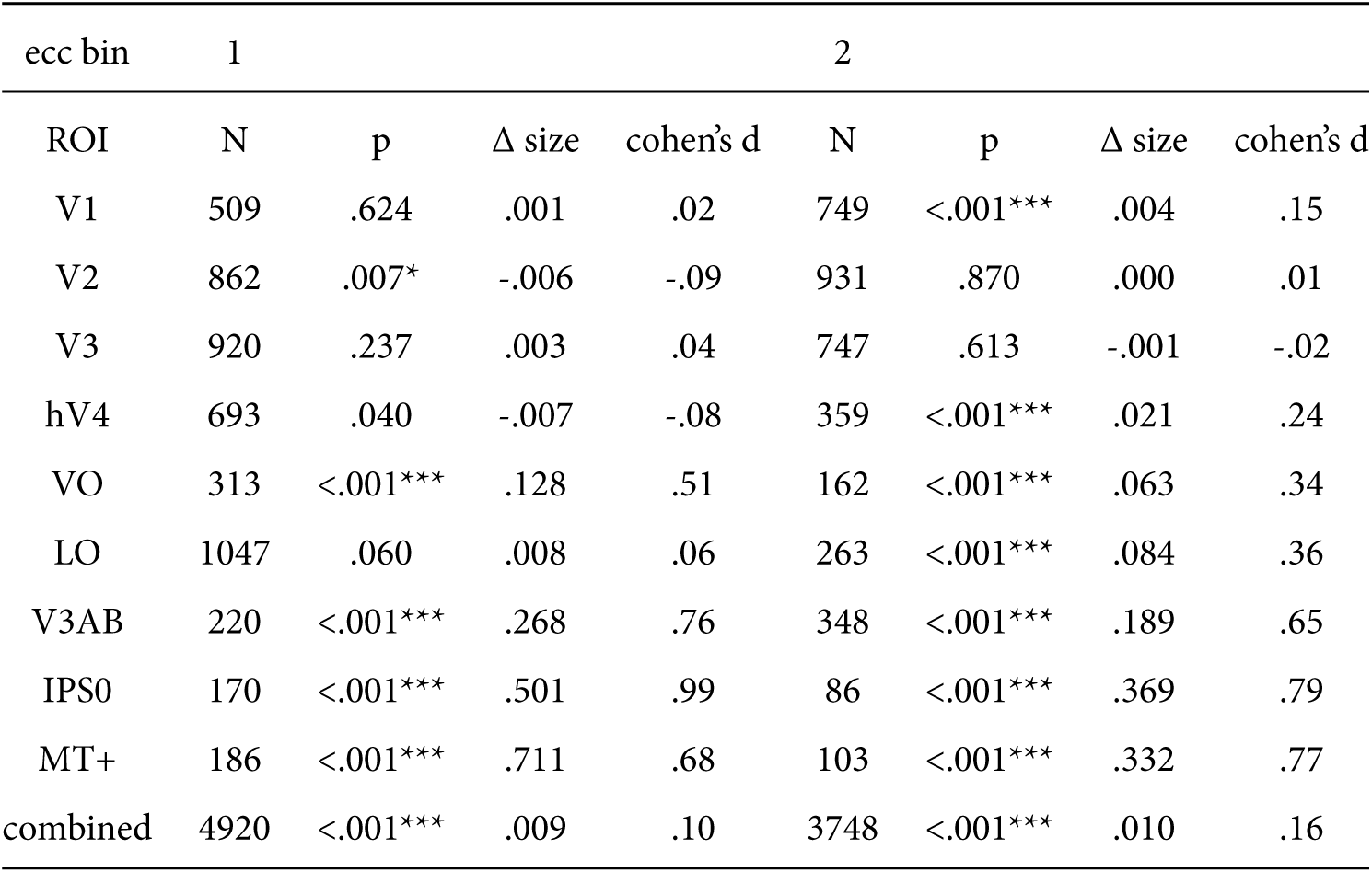
Statistics corresponding to Figure 3B on pRF size changes. P-values reflect whether bootstrapped pRF size difference distribution is different from 0, for each ROI and eccentricity bins 1 and 2 (bin 3 and 4 in table below). Single, double and triple asterisks indicate FDR corrected significance of <.05, <.01 and <.001 respectively. FDR correction performed over all p-values in this table simultaneously.

**Table 6:** Statistics corresponding to Figure 3B on pRF size changes. P-values reflect whether bootstrapped pRF size difference distribution is different from 0, for each ROI and eccentricity bins 3 and 4 (bin 1 and 2 in table above). Single, double and triple asterisks indicate FDR corrected significance of <.05, <.01 and <.001 respectively. FDR correction performed over all p-values in this table simultaneously.

| ecc bin | 3 |  |  |  | 4 |  |  |  |
| --- | --- | --- | --- | --- | --- | --- | --- | --- |
| ROI | N | p | $\Delta$ size | cohen's d | N | p | $\Delta$ size | cohen's d |
| V1 | 595 | <.001*** | .010 | .33 | 323 | .663 | .001 | .03 |
| V2 | 616 | .012* | -.005 | -.10 | 343 | <.001*** | -.021 | -.31 |
| V3 | 427 | <.001*** | -.015 | -.19 | 226 | <.001*** | -.052 | -.54 |
| hV4 | 110 | .012* | .029 | .24 | 39 | <.001*** | -.114 | -.72 |
| VO | 71 | <.001*** | .066 | .41 | 36 | <.001*** | -.085 | -.98 |
| LO | 56 | .108 | -.026 | -.21 | 31 | .006** | -.119 | -.45 |
| V3AB | 199 | <.001*** | .075 | .37 | 113 | <.001*** | -.111 | -.62 |
| IPS0 | 36 | <.001*** | .260 | .78 | 21 | .988 | .003 | .01 |
| MT+ | 18 | .541 | .056 | .14 | 18 | .128 | .227 | .31 |
| combined | 2128 | .252 | -.002 | -.03 | 1150 | <.001*** | -.035 | -.38 |

**Table 7:**
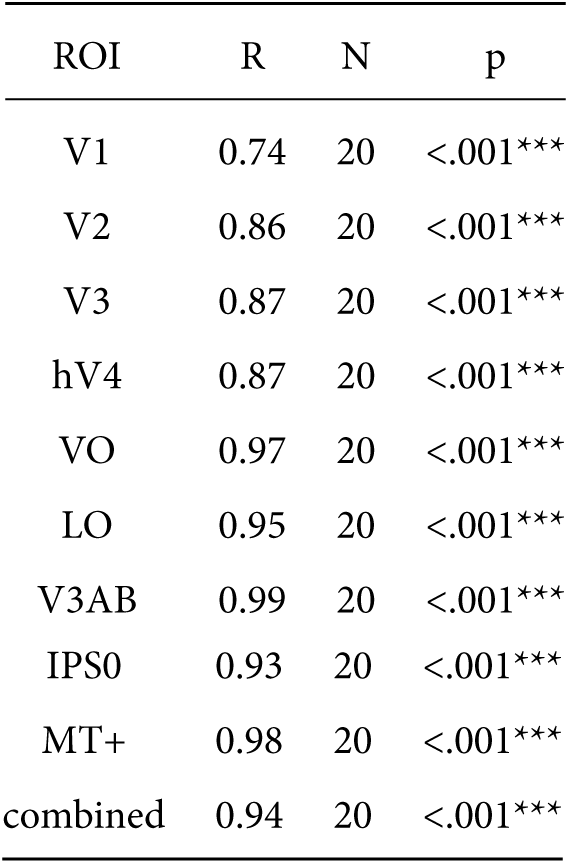
Statistics corresponding to Figure 3C on correlations between eccentricity and size changes. P-values are two-tailed tests whether bootstrapped distribution of pRF eccentricity and size change correlations across bins is different from 0. Triple asterisks indicate FDR corrected significance of <.001. FDR correction performed over all p-values in this table simultaneously.

In order to quantify the modulation of feature-based attention per voxel we first setup a single robust index of the of the degree to which spatial attention resampled visual space, combining changes in pRF eccentricity and size (as these were highly correlated, see Figure 3C). This Attentional Modulation Index (AMI, see Methods) is depicted in Figure 5B for the *combined ROI* when attending color and TF. We then quantified the difference in this attentional modulation index between attending color and temporal frequency as a feature-based Attentional Modulation Index (feature AMI, see Methods). Positive values of feature AMI indicate that attending color induced greater pRF changes, while negative values indicate that attending TF lead to stronger pRF changes. Figure 5C shows that this feature AMI was positive across eccentricity in the *combined ROI*. Inspecting the average feature AMI across voxels within each ROI (Figure 5D) reveals that attending changes in color compared to TF in the bar stimulus produced stronger spatial resampling in all ROIs (p’s < .01, see Supplementary Tables 8 and 23).

#### Feature AMI and feature preference

The feature-based modulations we describe above are possibly related to differences in bottom-up preference for the attended features. Feature-based attention is known to increase activity of neurons selective for the attended feature, regardless of spatial location (Treue and Maunsell, 1996; Treue and Trujillo, 1999; McAdams and Maunsell, 2000; Maunsell and Treue, 2006; Müller et al., 2006; Zhang and Luck, 2009; Zhou and Desimone, 2011). If voxels contain on average more color-preferring compared to TF-preferring neurons, attending color should therefore activate a greater amount of neurons, potentially leading to a greater apparent shift of the aggregate pRF. To test this hypothesis, we estimated the difference in response amplitude to the presence of color and temporal frequency within a full-field stimulus (in a separate experiment, see Methods). We then summarized each voxels’ relative preference for color and temporal frequency by means of a feature preference index. Higher values of this feature preference index indicate greater preference for color compared to TE Figure 5E (and Supplementary Figure 10) displays the feature AMI as a function of feature preference, for each ROI. Note that feature preference was negative on average in most ROIs, suggesting that our TF manipulation (7 vs 0 Hz grayscale Gabors) caused stronger response modulations compared to our color manipulation (colored vs grayscale Gabors). Although this induced an offset across the brain, variations in this measure across ROIs replicate known specializations of the visual system with high precision (Liu and Wandell, 2005; Brouwer and Heeger, 2009, 2013): while areas MT+ and V1 show the strongest preference for TF compared to color, areas V4 and VO show the strongest preference for color compared to TF. Importantly, regardless of these large variations in feature preferences between MT+/V1 and VO/hV4, average feature AMI was nearly equal in these ROIs. In addition, there was no correlation between feature preference and feature AMI across all ROIs (R = .20, p = .608, N = 9, rho = .37, p = .332, N = 9). These results show that the feature-based attentional modulations we observe occor globally across the brain, and do not depend on bottom-up feature preference.

#### Feature preference and spatial sampling

What could then explain the fact that attending color in the stimulus induced greater changes in spatial sampling? Behavioral studies have suggested that the influence of spatial attention should be adjusted by feature-based attention in order to improve sampling of attended visual features (Yeshurun et al., 2008; Barbot and Carrasco, 2017). One of the factors that influences required spatial resampling is pRF size. Smaller pRFs need to shift a greater distance in order to bring a stimulus into their responsive region. This means that if color-prefering voxels are relatively small, this creates a requirement of greater shifts when attending color. Indeed, both pRF size (Dumoulin and Wandell, 2008) and color compared to TF preference (Curcio et al., 1990; Azzopardi et al., 1999; Brewer et al., 2005) are known to be strongly eccentricity dependent such that foveal voxels have relatively small pRFs and are relatively color sensitive. We also clearly observe both effects in our data (see Figure 1B and Supplementary Figure 1; and Figure 6 and Supplementary Figure 11, correlation between feature-preference and eccentricity is negative except in LO and VO, see Supplementary Tables 9 and 24). This means that the greater amount of spatial resampling when attending color can be parsimoniously explained by color being sampled by relatively smaller pRFs.

**Figure 6:**
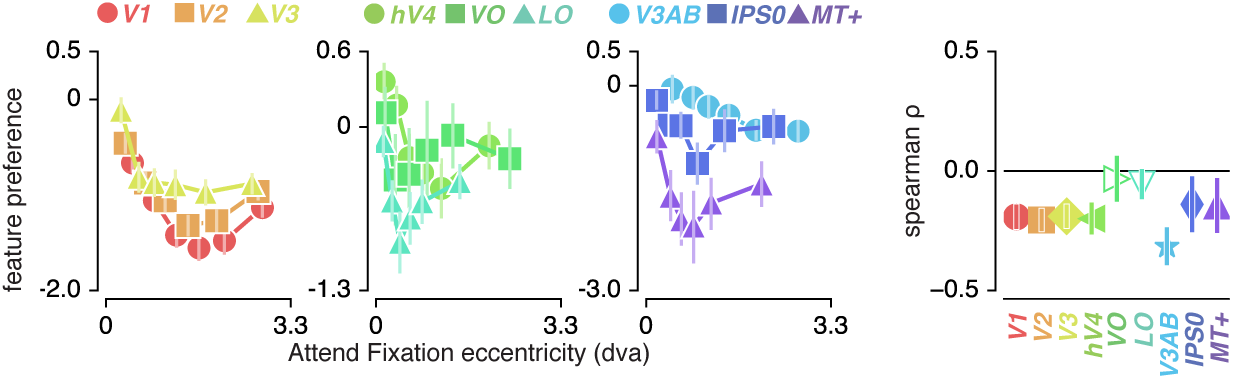
Feature preference and eccentricity. Preference to color compared to TF is greatest near the fovea. Errorbars denote 95% CI over voxels.

#### Feature-based attention influences attentional gain field precision

Smaller pRFs also require a more precise attentional gain field to shift a given distance (a property of the multiplication of Gaussians). Combining this with our observation that pRFs experience greater shifts when attending color, we predict that attentional gain fields should be more precise in this condition. In order to test this, we repeated the attentional gain field modeling procedure described above, replacing the *Attend Stimulus* data with the *Attend Color* and *Attend TF* data in two seperate fit procedures. Indeed, this returned smaller fitted stimulus attentional gain field sizes in the *Attend Color* compared to the *Attend TF* fit procedure (Figure 7 left panel) both in the *combined ROI* (0.094 dva smaller over subjects when attending color, t(4) = 9.021, p = .001, cohen’s d = 4.511) and across ROIs (median over ROIs on average 0.061 dva smaller over subjects when attending color, t(4) = 4.243, p = .013, cohen’s d = 2.121). As the *Attend Fixation* data was used as input in both modeling procedures, we verified that the estimated fixation attentional gain field was not different between procedures (Figure 7 right panel; p’s of .693 and .224 and cohen’s d of −0.213 and −0.719 for across ROIs and *combined* ROI respectively). These analyses show that the stronger influence of spatial attention when attending color is realized by a more precise attentional gain field located at the stimulus. In sum, our results suggest that (1) the attentional system adjusts its influence in accordance with the spatial sampling characteristics of units that prefer the attended feature and (2) that it does this equally across visual regions regardless of their bottom-up feature preference.

**Figure 7:**
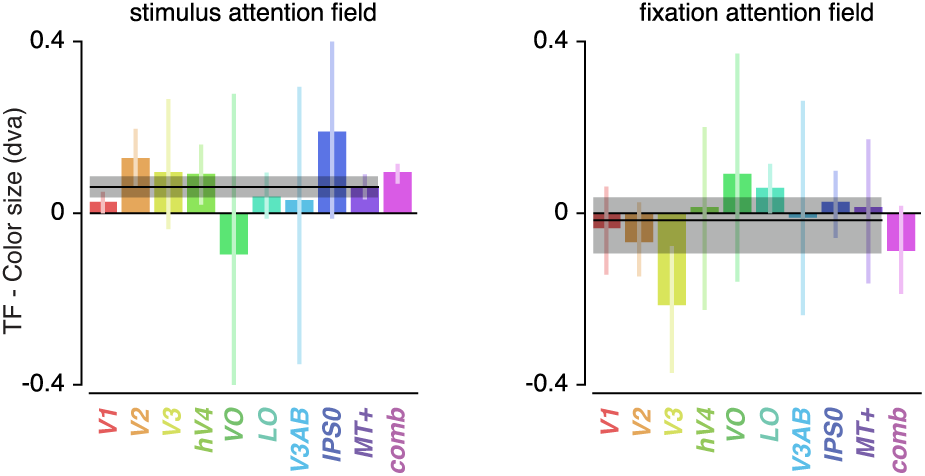
Feature-based attentional modulation of attentional gain field sizes. The greater effects of attention to color compared to TF are caused by more precise attentional gain fields toward the stimulus. Error bars represent 95% CI over subjects. Black horizontal lines indicates median attentional gain field size difference over ROIs, with shaded area representing 95% CI over subjects.

### Task and Fixation Performance

Finally, we verified that the pRF results were not affected by differences in fixation accuracy or behavioral performance (see Figure 8). To provide evidence in favor of these null hypothesis, we performed JZL Bayes factor analyses (using JASP; Love et al. (2015)). We rotated recorded eye position to the direction of bar movement and computed the median and standard deviation of position along this dimension across bar passes per bar position (Figure 8B). We next setup a model including the factor of attention condition (3 levels), bar position (24 levels), and their interaction. We found that when predicting gaze position, the evidence was in favor of the null hypothesis with a Bayes Factor (BF) of 18620. When predicting gaze variability however, we found evidence against the null hypothesis with a BF of 5.980. Evidence for including each of the factors (condition, bar position, and their interaction) into the model returned BFs of 0.713, 547.193 and 0.017 respectively. The BF of 0.713 for the factor of condition means that we cannot determine whether gaze variability was different between conditions. However, even if this would be the case, this could only lead to an offset in pRF size and not to changes in pRF position (Levin et al., 2010; Klein et al., 2014; Hummer et al., 2016). As we observe a complex pattern of both pRF size and eccentricity in- and decreases, a difference in gaze variability cannot explain the attentional modulations of pRF parameters we observed. More importantly, these analyses also showed that although bar position influenced gaze variability (BF of 547.193), it did not do so differently between attention conditions (BF of 0.017). Using a similar approach, we then tested whether a model including attention condition (3 levels) and stimulus eccentricity (3 levels) influenced behavioral performance (Figure 8A). This returned evidence for the null hypothesis with a BF of 6.25. Together, these results show that differences in pRF parameters between conditions cannot be explained by either fixation accuracy or behavioral difficulty.

**Figure 8:**
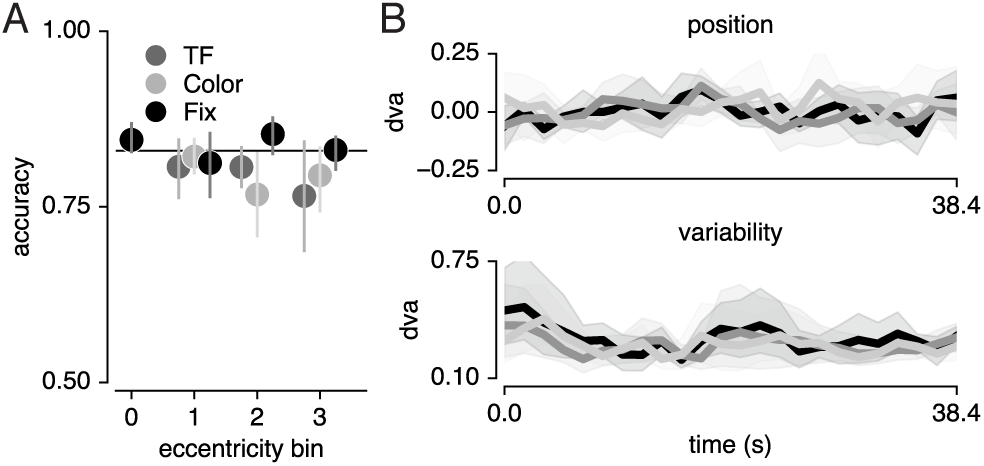
Task and fixation performance A. Behavioral accuracy per attention condition and per bar stimulus eccentricity bin. Horizontal line denotes Quest target of 83%; chance level was 50%. B. Median (top pannel) and standard deviation (bottom pannel) gaze position in the direction of bar movement per bar position. Error bars denote 95% CI across 5 participants.

## Discussion

We investigated how spatial and feature-based attention jointly modulate the sampling of visual space. We find that directing covert spatial attention toward a moving bar stimulus altered the eccentricity and size of pRFs in concert. These changes in spatial sampling were parsimoniously explained by an attentional gain field model. Attending color changes within this stimulus induced stronger pRF changes compared to attending temporal frequency changes. These feature-based attentional modulations occured globally throughout the brain, irrespective of a visual region’s average feature preference. We suggest that the greater degree of spatial resampling when attending color is related to smaller pRF sizes in relatively color preferring voxels. In addition, we show that the greater degree of spatial resampling when attending color is caused by a more precise attentional gain field on the stimulus.

The average changes in pRF size and eccentricity for each visual region are largely consistent with previous studies where attention was devoted to a peripheral stimulus versus fixation (Kay et al., 2015; Sheremata and Silver, 2015). More specifically, we systematically investigated the spatial structure of the complex pattern of pRF changes that resulted from such differential spatial attention. The resulting characterization details how the sampling of visual space by single voxel pRFs is affected by spatial attention, which is of specific relevance for future studies that determine spatial selectivity for voxel selections. First, we show that attending the stimulus compared to fixation caused pRFs to shift radially. Although a previous study reported a dominant horizontal shift direction (Sheremata and Silver, 2015), we suggest that the overrepresentation of the horizontal meridian (Schneider et al., 2004; Swisher et al., 2007) made radially shifting pRFs to appear as predominantly horizontal changes. Second, we report closely coupled pRF eccentricity and size changes that depended on pRF eccentricity. Specifically, we found that parafoveal pRFs shifted toward the periphery and increased in size while peripheral pRFs shifted toward the fovea and decreased in size. This finding supports the resolution hypothesis of attention (Anton-Erxleben and Carrasco, 2013), which posits that spatial attention acts to reduce resolution differences between the fovea and periphery. We note that the functional implication of pRF size changes was recently questioned, as stimulus encoding fidelity was shown to be unaffected by pRF size changes (Vo et al., 2017). However, the exact functional significance of changes in pRF size bears no consequence for the conclusions currently presented. Due to the strong correlation we observed between pRF eccentricity and size changes, we combined both measures into a single robust index. Our relevant quantifications are therefore agnostic to the potentially separable functional implications of changes in pRF size and eccentricity.

The pattern of pRF shifts we observe is well described by an attentional gain field model (Reynolds and Heeger, 2009; Klein et al., 2014). First, this highlights that a simple and well-understood mechanism underpins the apparent complexity of the observed pattern of pRF changes. Second, it extends the utility of such attentional gain field models to situations where attention is dynamically deployed over space and time during the mapping of the pRF (Kay et al., 2015). In agreement with earlier reports (Klein et al., 2014; Puckett and DeYoe, 2015), we found that the best-fitting model implemented comparable attentional gain field sizes across visual regions. This strongly points to spatial attention being implemented as a global influence across visual cortex. We conclude that differences in pRF shift patterns between different visual regions depended primarily on differences in visual sampling (i.e. differences in pRF center and size distributions) rather than on differing attentional influences. Despite the broad correspondence between model fits and data, the model did not capture the observed decreases in pRF eccentricity of the most foveal pRFs in VI. Two recent studies showed that in early visual areas, spatial attention shifted pRFs away from the attended location, but toward the attended location in higher visual areas (de Haas et al., 2014; Vo et al., 2017). Other studies showed that in precisely these visual regions, both the pRF and the attentional gain field are composed of a suppressive surround in addition to their positive peak (Zuiderbaan et al., 2012; Puckett and DeYoe, 2015). We leave the question whether these suppressive surrounds could explain such repulsive shifts in lower visual cortex for future research. As a more general aside, gain fields have been shown to influence visual processing at the motor stage (van Opstal et al., 1995; Snyder et al., 1998; Trotter and Celebrini, 1999). Thus, our results further establish the close link between attentional and motor processes (Rizzolatti et al., 1987; Corbetta et al., 1998).

Of particular interest, we showed that the changes in visual space induced by spatial attention were stronger when attending color compared to when attending temporal frequency. This occured throughout the brain, irrespective of visual regions’ preference for the attended features. In other words, while MT+ and V4 differed greatly in their relative feature preference, both areas showed comparable pRF changes resulting from differences in feature-based attention. This stands in apparent contrast with previous studies reporting that feature-based attention selectively enhances responses in cortical areas specialized in processing the attended feature (Corbetta et al., 1990; Chawla et al., 1999; O’Craven et al., 1999; Schoenfeld et al., 2007; Baldauf and Desimone, 2014). However, attending a stimulus consisting multiple features was shown to spread attentional response modulations of one of the object’s feature dimensions to other constituent feature dimensions (Katzner et al., 2009; Çukur et al., 2013; Kay and Yeatman, 2017), albeit somewhat later in time (+/− 60ms; Schoenfeld et al., 2014). This could mean that the global pattern of pRF shifts we observed is caused by such an object-based attentional transfer mechanism. In addition, changing the sampling of visual space globally throughout the brain enhances stability in the representation of space. Different modifications of visual space per visual region would require an additional mechanism linking different spatial representations. Instead, the global nature of spatial resampling we observe supports a temporally dynamic but spatially consistent adaptation of visual space.

The greater degree of spatial resampling when attending color can be explained by differences in spatial sampling in voxels that prefer color compared to temporal frequency. Behavioral studies have suggested that feature-based attention should modulate the strength of spatial resampling in order to improve sampling of attended features (Yeshurun et al., 2008; Barbot and Carrasco, 2017). One of the factors that determines the required degree of spatial resampling is receptive field size. Specifically, smaller pRFs need to shift a greater distance in order to bring a stimulus into their responsive region. Both our data and previous findings show that pRF size (Dumoulin and Wandell, 2008) and color compared to temporal frequency preference (Curcio et al., 1990; Azzopardi et al., 1999; Brewer et al., 2005) vary across eccentricity such that foveal voxels have smaller pRFs and are more color sensitive. This means that color is sampled by smaller pRFs than temporal frequency. Attending color in the stimulus therefore required a greater degree of spatial resampling in order to optimally sample the attended feature. In addition, gain field models of attention predict that smaller pRFs require a more precise attentional gain field in order to shift a given distance. Indeed, our results showed that attending color versus temporal frequency resulted in more precise attentional gain fields directed toward the bar stimulus. Together, this shows that the larger degree of spatial resampling we observed when attending color can be parsimoniously explained by smaller pRF size in voxels that prefer color over temporal frequency. Yet, as pRF size is so strongly related to pRF eccentricity, we can not exclude an influence of pRF eccentricity above and beyond pRF size. However, receptive field size is suggested to be crucial in the eccentricity dependence of attentional influences observed in behavior (Yeshurun and Carrasco, 1998, 1999; Anton-Erxleben and Carrasco, 2013). Therefore, although differential influences of attention across eccentricities have been observed before in the brain (e.g. Roberts et al., 2007; Bressler et al., 2013), these are likely brought about by differences in pRF size. In sum, our results confirm the notion that the endogenous attentional system is able to take into account the spatial sampling properties of units that prefer the attended feature, and that it adjusts the strength of its influence accordingly (Yeshurun et al., 2008; Barbot and Carrasco, 2017).

Electrophysiological studies on the interaction between feature-based and spatial attention generally measure overall response amplitudes instead of changes in spatial sampling. This showed that interactions between feature-based and spatial attention are weak to non-existent in relatively early stages of processing (David et al., 2008; Hayden and Gallant, 2009; Patzwahl and Treue, 2009; Zhang and Luck, 2009), but develop at later stages of visual processing (Hillyard and Munte, 1984; Handy et al., 2001; Andersen et al., 2011; Bengson et al., 2012; Ibos and Freedman, 2016, but see Egner et al. (2008)). We add to this (1) that feature-based attention modulates the effects of spatial attention on spatial resampling and (2) that these interactions occur globally throughout the brain, manifesting themselves even in the earliest cortical visual regions. Interactions between spatial and feature-based attention in early stages of processing could be concealed when focusing on changes in response amplitude instead of changes in spatial sampling. However, it is important to note that measuring spatial sampling at the level of voxels does not allow us to determine whether observed changes in spatial sampling are due to changes in spatial sampling of individual neurons, or rather due to differential weighting of subpopulations of neurons within a voxel. Nevertheless, it has been shown that spatial attention does influence spatial sampling of individual neurons (Connor et al., 1997; Womelsdorf et al., 2006). Yet, future studies are required to extend our conclusions regarding the interactions between feature-based and spatial attention to the single neuron level.

Another important remaining question pertains to the source of the interactions between feature-based and spatial attention. Signals of spatial selection are thought to originate from a network of frontal and parietal areas, identified using fMRI (Shulman, 2002; Silver et al., 2005; Jerde et al., 2012; Sprague and Serences, 2013; Szczepanski et al., 2013; Kay and Yeatman, 2017; Mackey et al., 2017), and electrophysiology (Moore and Armstrong, 2003; Gregoriou et al., 2009). As we focused on careful measurement of spatial sampling in feature sensitive visual cortex with a relatively small stimulus region, we did not include the frontoparietal regions containing large receptive fields into our analyses. A recent study suggested a central role for the ventral prearcuate gyrus for conjoined spatial and feature-based attentional modulations (Bichot et al., 2015). Correspondingly, signals of feature selection in humans have been localized to a likely human homologue of this area, the inferior frontal junction (IFJ; Zanto et al., 2010; Baldauf and Desimone, 2014). This region is therefore a possible candidate for controlling the interactions between feature-based and spatial attention.

In sum, we show that visuospatial sampling is not only affected by attended locations but also depends on the spatial sampling properties of units that prefer attended visual features. The global nature of these modulations highlights the flexibility of the brain’s encoding of sensory information in order to meet task demands (Rosenholtz, 2016).

## Materials and Methods

### Participants

Five participants (2 female, 2 authors, aged between 25 - 37) participated in the study. All gave informed consent, and procedures were approved by the ethical review board of the University of Amsterdam, where scanning took place.

### Apparatus

#### MRI acquisition

All MRI data was acquired on a Philips Achieva 3T scanner (Philips Medical Systems), equipped with a 32-channel head coil. T1 weighted images were acquired for each participant with isotropic resolution of 1 mm^3^, repetition time (TR) of 8000 ms, TE of 3.73 ms, flip angle of 8°. Functional T2* weighted data consisted of 30 2D slices of echo planar images (EPI) with isotropic resolution of 2.5 mm^2^, with a 0.25 mm slice gap, TR of 1600 ms, TE of 27.62 ms, and a flip angle of 70°. Each participant completed between 6 to 8 Attention-pRF Mapping runs (20 min each) and 2-3 Feature preference and HRF Mapping runs (10 min each), spread over 2 (N = 1) or 3 (N = 4) sessions within a 2 week period (see Experimental Design).

#### Gaze recording

During all functional runs, gaze position was recorded using an Eyelink 1000 (SR Research, Osgoode, Ontario, Canada), sampled at 1000 Hz. A 9-point calibration-validation procedure was run at the start of each session.

#### Stimulus presentation

Visual stimuli were created in PsychoPy (Peirce, 2008) and run on a 15 inch 2013 MacBook Pro Retina. Participants viewed a 32 inch BOLD screen (resolution: 1920 x 1080, refresh rate: 100 Hz; Cambridge Research Systems), at 156 cm distance of the participants’ eyes at the end of the bore, via a helmet-mounted front-silvered mirror. Auditory stimuli were presented through headphones using the MRConfon system.

### Experimental Design

#### Attention-PRF Mapping Stimulus

A bar stimulus of 0.9 degrees of visual angle (dva) width traversed a circular aperture of 7.2 dva in one of eight directions (cardinal and diagonal orientations in both directions, see Figure 9A), completing a full pass in 38.4 s by stepping 0.34 dva every 1.6 s, and pausing 4.8 s between each bar pass. One run contained 24 bar passes in total (3 for every direction), plus four blank periods of 38.4 s when no bar stimulus was shown. Throughout the experiment, a gray disk of 9.6 arcmin (60 cd/m^2^), with a 4.2 arcmin black rim (0 cd/m^2^) was present on the screen as a fixation mark. The bar stimulus was composed of 1500 Gabor elements (4.34 cycle/dva spatial frequency, 9 arcmin sd, average luminance of 60 cd/m^2^) projected over a dark-gray background (15 cd/m^2^). Three times per bar location (i.e. every 533 ms), Gabor element parameters were updated to a new random location (uniformly distributed over the spatial extent of the bar at full width), a random orientation (uniformly drawn between 0 - 360°), a random color combination (either blue-yellow (BY), or cyan-magenta (CM)) and a random new temporal frequency (TF; either high or low). The high and low temporal frequencies were chosen per participant to facilitate their ability to distinguish TF changes (6 and 4 Hz in 3 participants, 7 and 3 Hz in 2 participants). The overal color and/or TF composition of the bar was transiently altered on some of these parameter updates, by changing the ratio of Gabor elements assigned either color combination or either TF (as targets for the behavioral tasks, see below). The temporal predictability of these events was minimized by randomly drawing occurences according to an exponential distribution (mean 4 s, minimum 2 s). Additionally, the fixation mark central disk luminance either increased or decreased, with probability and duration of these occurrences identical to those of changes in the bar stimulus composition. These three types of transients (fixation mark luminance, bar color and TF composition) were independent, meaning they were randomly interleaved and could be combined on the screen. Importantly, this design ensured that physical stimulation was equated across all three attention conditions, which we describe below.

**Figure 9:**
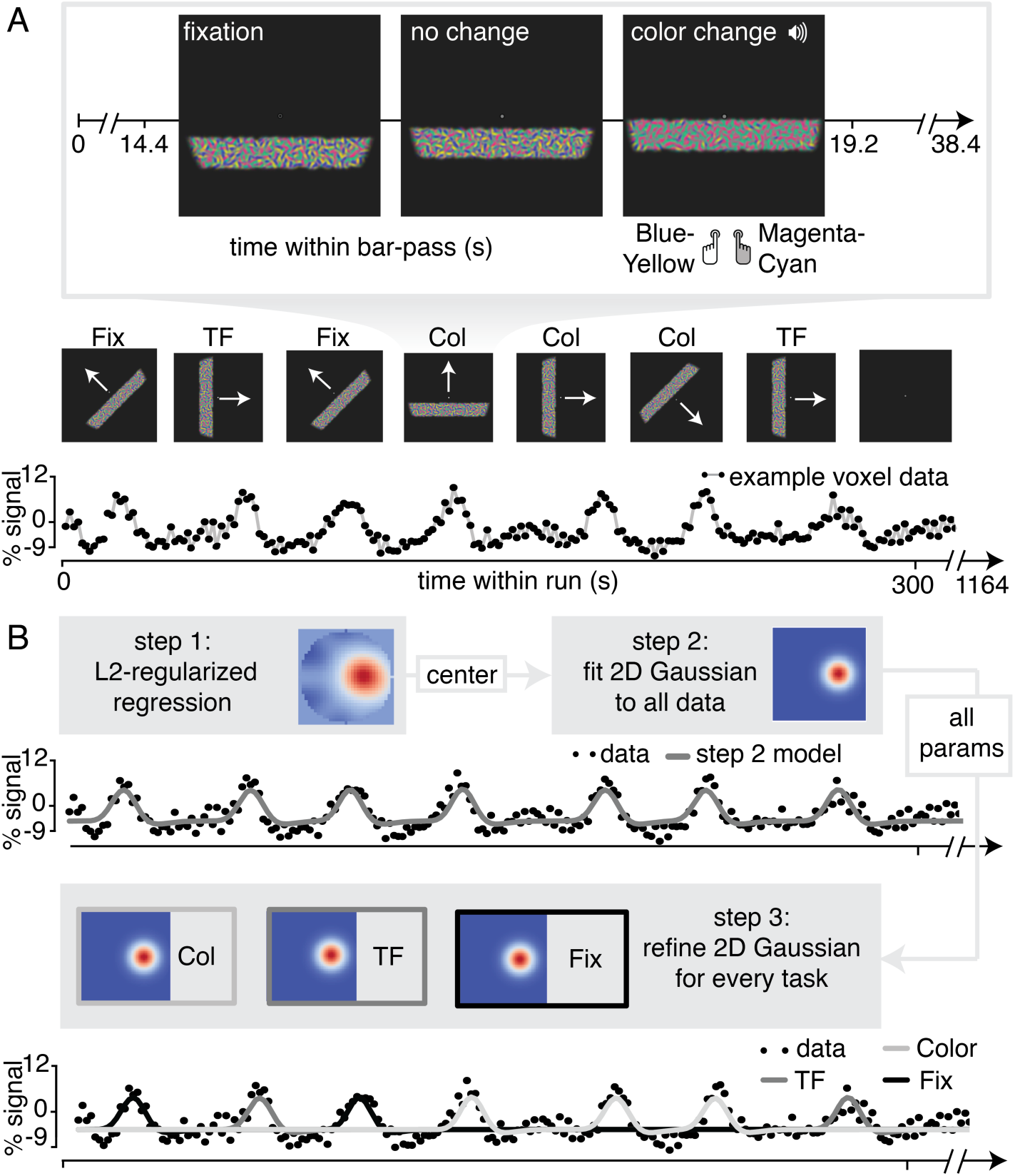
Experimental Design and pRF Fitting procedure. A. Experimental Design. Throughout a bar pass participants reported either changes in color (*Attend Color*) or temporal frequency (*Attend TF*) of Gabor elements within the moving bar stimulus, or changes in fixation mark luminance (*Attend Fixation*), while maintaining accurate fixation. Participants were informed auditorily about the upcoming task 2 s before each bar pass. B. Overview of pRF fitting procedure. pRF parameters were estimated from each voxel’s BOLD signal time course in a three step procedure. First, a design matrix was constructed based on 31×31 pixels’ visual stimulation time course of the entire experiment, which was convolved with a participant-specific HRF (derived from separate data, see *Feature preference and HRF Mapper*). L2-regularized regression was used to find the position of the spatial response profile’s peak of each voxel. Second, to find precise estimates of pRF center location and size we used a more finegrained 101 × 101 design matrix and gradient descent to fit a single parameterized 2D Gaussian pRF model to data from all attention conditions combined, initialized at the L2-regression derived peak location. Third, 2D Gaussian pRF models were fitted to data from the different attention conditions separately, intialized with the parameters resulting from step 2.

#### Attention-PRF Mapping Task

For an overview of the stimulus and behavioral task, see Figure 9A. Before each bar pass an automated voice (Apple OSX Dictation voice ‘Kathy’) informed participants to perform a 2-alternative forced-choice task (2AFC) on one of the three stimulus parameter deviations. Task-relevant visual stimulus changes were accompanied by an auditory pure tone (440 Hz). This auditory cue alerted the participant to respond, while task-irrelevant stimulus changes occurred independently and without warning tone. This ensured that all task-related information was conveyed to the participant by auditory means, without concurrent changes in visual stimulation. The different stimulus changes (i.e. color, TF and fixation luminance) occurred independently and thus sometimes simultaneously, rendering the auditory tone not reliably predictive of the stimulus dimension to attend. Therefore, participants needed to stably maintain condition-specific top-down attention throughout the duration of a bar pass. In the *Attend Color* condition, participants judged the relative predominance of Blue-Yellow or Cyan-Magenta Gabor elements in the bar stimulus, while in the *Attend TF* condition, participants judged the relative predominance of high compared to low TF Gabor elements in the bar stimulus. We chose color and TF as these features are known to be processed markedly differently in the visual system. While color is preferentially processed by the ventral visual areas V4 and VO, temporal information is known to be processed preferentially by area MT+ (Liu and Wandell, 2005; Brouwer and Heeger, 2009, 2013). We specifically chose TF and not coherent motion, as coherent motion signals have been shown to influence pRF measurements (Harvey and Dumoulin, 2016). Both color and TF have been shown to be able to capture attention (Wolfe and Horowitz, 2004; Cass et al., 2011). In the *Attend Fixation* condition, participants judged whether the central disk of the fixation mark increased or decreased in luminance. The magnitude of the stimulus changes was titrated by means of a Quest staircase procedure (Watson and Pelli, 1983), set to approximate 83% correct performance. In order to equate task difficulty across not only conditions but also bar stimulus eccentricity, we used separate Quest staircases at three different bar stimulus eccentricies in each of the attention conditions. Additionally, there was a separate staircase for the *Attend Fixation* task when no bar stimulus on screen. This made for a total of 10 separate staircases during the experiment. Participants extensively practiced the task outside the scanner and staircases were reset before scanning. Each experimental run contained one bar pass per task condition, per direction, in random order (total of 24 bar passes per run).

#### Feature preference and HRF Mapper

We performed a separate randomized fast event-related fMRI experiment in order to (1) determine each voxels’ relative preference for color and TF, and (2) to find the parameters that best described each participants’ HRF, to be used in the pRF estimation procedure (see below). Full-field stimuli consisted of 8000 Gabor elements, uniformly distributed throughout the full circular aperture traversed by the pRF mapping stimulus ensuring identical density compared to the pRF mapping stimulus. Also, every 533 ms, all Gabor elements were assigned a new random orientation and location. These stimuli were presented for 3.2 s, with an inter-trial interval of 3.2 s. In a full factorial 2 × 2 design, we varied the color and TF content of the stimulus in an on-off fashion. That is, the TF of the Gabor elements was either 0 or 7 Hz, and the elements were either grayscale or colored (balanced BY/CM). Trial order was determined based on an M-sequence (Buračas and Boynton, 2002), with no-stimulus (null) trials interspersed as a fifth category of trials. During this experiment, participants performed the same 2-AFC fixation-point luminance task as in the Attention-pRF Mapping Task (*Attend Fixation*), using a separate staircase. A single HRF was determined per participant using the Rl-GLM approach (Pedregosa et al., 2015), on data from all conditions. The median HRF from the 1000 most responsive voxels (highest beta-weights in the colored high TF condition) was used as the participant specific HRF.

### Data analysis

#### MRI Preprocessing

T1-weighted images were first segmented automatically using Freesurfer, after which the pial and grey/white matter surfaces were hand-edited. Regions of interest (ROIs) were defined on surface projected retinotopic maps using Freesurfer without the use of spatial smoothing. For every participant, one session’s EPI image was selected as the target EPI, which was registered to his/her Freesurfer segmented T1-weighted image using the bbregister utility, after which the registration was hand-adjusted. Then, all EPI images were first motion corrected to their middle volume using FSL (Jenkinson et al., 2012) MCFLIRT to correct for within run motion. Then, all EPI images were registered both linearly (using FLIRT) and non-linearly (using FNIRT) to the mean-motion corrected target EPI to correct for between run and session motion and inhomogeneities in B0 field. Low frequency drifts were removed using a 3rd order savitzky-golay filter (Savitzky and Golay, 1964) with a window length of 120s. Arbitrary BOLD units were converted to percent-signal change on a per-run basis.

#### pRF fitting procedure

pRF fitting and (statistical) parameter processing was performed using custom-written python pipeline available at https://github.com/daanvanes/PRF_attention_analyses. Links to the data files required to reproduce the figures and analyses can be found in the readme of this repository. The fitting routines heavily relied on the scipy and numpy packages. We approximated the pRF by a two-dimensional isotropic Gaussian function. For an overview of our pRF fitting procedure, see Figure 9B. A predicted timecourse for a given Gaussian function can be created by first computing the overlap of this function with a model of the stimulus for each timepoint, and then convolving this overlap with the participant-specific HRF (Dumoulin and Wandell, 2008). It is possible to find these Gaussian parameter estimates using a minimization algorithm, but such an approach is at risk of missing the global optimum when parameters are not initialized at appropriate values. Recently, a model-free reverse-correlation-like method was developed, generating a pRF spatial profile without requiring any pre-set parameters (for details see Lee et al. (2013)). Briefly, we employed this method using L2 regularized (Ridge) regression on a participant-specific-HRF convolved design matrix coding the stimulus position in a 31×31 grid for each timepoint, predicting data from all attention conditions together. Using a high regularisation parameter (*λ* = 10^6^), we used this procedure not to maximize explained signal variance, but to robustly determine the pRF center, which was defined as the position of the maximum weight. Having determined these approximate initial values for the pRF center, we next initialized a minimization procedure (Powell (1964) algorithm) at these location values, fitting position (x, y), size, baseline and amplitude parameters of an isotropic 2D Gaussian to data from all conditions together using a design matrix with size 101×101 for enhanced precision. Then, all resulting Gaussian parameters were used to initialize a second minimization procedure which fitted a Gaussian for each attention condition separately at the same time (all parameters independent except for one shared baseline parameter). This approach allowed us to recover fine-grained differences in pRF parameters under conditions of differential attention.

#### pRF selection

We discarded pRFs that were either at the edge of the stimulus region (above 3.3 dva in the *Attend Fixation* condition), or had size (standard deviation) larger than our stimulus diameter (7.2 dva) in any of the tasks. Additionally, each voxel’s contribution to all analyses was weighted according to the quality of fit of the pRF model, which was defined as 1 minus the ratio of residual to observed variance:

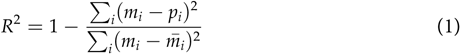

where *i*, *m* and *p* refer to voxel index, measured BOLD time-course and predicted time-course respectively. We disregarded voxels with an *R^2^ <* .1.

#### pRF parameter analyses

The statistical approach used to combine pRF results across participants was similar to that of two recent and comparable studies (Klein et al., 2014; Kay et al., 2015), that adopted a ‘dense sampling of individual brain approach’ (Poldrack, 2017). This approach favours careful measurement of individual brains at the expense of large sample sizes in terms of the number of subjects. After defining visual ROIs per participant using standard retinotopic mapping procedures (Dumoulin and Wandell, 2008), this entailed pooling voxels for each ROI across participants. Statistics are then computed over this collection of voxels, much akin to the way in which neurons are often pooled across monkeys. In order to provide additional estimates of stability across participants, we also performed all analyses for each participant separately, the results of which can be found in the Statistical Appendix.

Except for the Rayleigh tests (Supplementary Table 2) p-values and confidence intervals were computed using 10^5^ fold bootstrap procedures. To test whether bootstrapped distributions differed from a certain threshold, p-values were defined as the ratio of boot-strap samples below versus above that threshold multiplied by 2 (all reported p-values are two-tailed). In order to provide maximally robust estimates of central tendency and variance of parameter estimates, we excluded outliers using a threshold of five two-sided median absolute deviations. As this outlier rejection was performed for each analysis separately, this resulted in slightly different number of voxels included in each analysis (see Ns in statistics tables). Outlier rejection was performed either per visual field bin (cf. Figure 1B, 2B/D, 3A/B, 4A/B/D/E/F, 5A/B, scatters in Figure 6), per percentile bin (cf. Figure 3C), or per ROI (cf. Figures 2C, 5D/E). When analyses are performed across participants outliers were again rejected at the voxel level per participant, but all participants were always included (cf. 4A/B/C/D/E/F and 7). When comparing correlations to 0, correlations were Fisher transformed using the inverse hyperbolic tangent function.

#### Feature attention modulation index

We computed a per-voxel index to quantify how strongly feature-based attention modulated the effects of spatial attention on spatial sampling (feature-based attention modulation index, or feature AMI). This measure combined pRF eccentricity and size parameters, as our results showed that spatial attention affected these parameters in concert (see Figure 3). Per voxel and per attention condition to the bar stimulus, (*Attend Color* and *Attend TF*) we set up a two-dimensional vector containing difference in pRF eccentricity and size relative to the *Attend Fixation* condition. pRF size and eccentricity differences were normalized respective to their variance in the *Attend Stimulus* condition to ensure that both measure contributed equally to the feature AMI. We then computed a feature AMI by dividing the difference between the norms of these vectors by their sum. This way, positive values of feature AMI indicate greater spatial attention effects on pRF parameters in the *Attend Color* condition than in the *Attend TF* condition and vice versa. Note that this measure abstracted out both the affected pRF parameter (i.e. eccentricity and size) and the sign of these changes (i.e. shifts toward the fovea or periphery and increases or decreases in size).

#### Attentional Gain Field modeling

To provide a parsimonious mechanistic account for how attention to the moving bar stimulus changed pRF position, we adapted an existing attentional gain model of attention (Womelsdorf et al., 2008; Reynolds and Heeger, 2009; Klein et al., 2014). This model conceptualizes the measured Gaussian pRF as the multiplication between a Gaussian stimulus-driven pRF (SD, i.e. the pRF outside the influence of attention), and a Gaussian attentional gain field (*AF*). Following the properties of Gaussian multiplication, the narrower the AF the stronger the influence on the SD, and the narrower the SD the smaller the resulting shift. An overview of model mechanics is shown in Figure 10. We estimated the SD by dividing the measured *Attend Fixation* pRF by an AF at fixation (*AF_fix_*, Figure 10A). As attention to the stimulus shifted the locus of attention as the bar moved across the screen, we modeled the effect of attention for each bar stimulus position separately (Figure 10B). Each unique bar stimulus (24 bar positions for each of 8 directions) was convolved with a Gaussian kernel (*AF_stim_*) and multiplied with the estimated SD. This yielded one predicted pRF per bar position. These predicted pRFs were then scaled to a maximum of 1 and averaged over the different bar positions. This averaged profile essentially represented the pRF ‘smeared’ across visual space as spatial attention moved along with the bar stimulus throughout the recording. The peak of this averaged profile was then designated as the predicted pRF center location in the *Attend Stimulus* condition (Figure 10C). Thus, the modeling procedure consisted of two separate AFs, one at fixation (*AF_fix_*) and one convolved with the bar stimulus (*AF_stim_*), which were estimated at the same time. The input to the model was the *Attend Fixation* pRF, and the output was the predicted position for the *Attend Stimulus* pRF. Formally, this is given by:

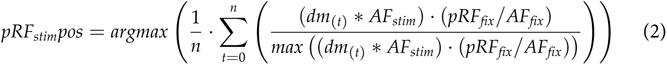

where (*dm*_(*t*)_ * *AF_stim_*) represents the stimulus design matrix at timepoint *t* convolved with the AF toward the stimulus, (*pRF_fix_/AF_fix_*) represents the estimation of the SD, and the denominator ensures scaling to a maximum of 1. To estimate how well a set of AF parameters fit the data across the entire visual field, we minimized the L2 distance between the predicted and measured *Attend Stimulus* pRF positions of the 64 vectors derived from the quadrant visual field format of Figure 2B. AF sizes were determined at an ROI level, thus assuming that attention influenced all pRFs within an ROI similarly, while possibly varying between ROIs. The model was evaluated for a 50 × 50 evenly spaced grid of AF sizes, where the AF at fixation varied between 1.5-2.5 dva, and the AF convolved with the stimulus varied between 0.6-1.6 dva (i.e. 0.02 dva precision). The convolution between the stimulus and the *AF_stim_* resulted in effective AF size to be 0.9 (bar stimulus width) larger than the *AF_stim_* itself. These parameter ranges therefore result in equal effective AF sizes. Reported sizes are the standard deviation of the 2D Gaussians, with 0.9 (the bar width) added to *AF_stim_* sizes. Modeling was performed for each participant separately.

**Figure 10:**
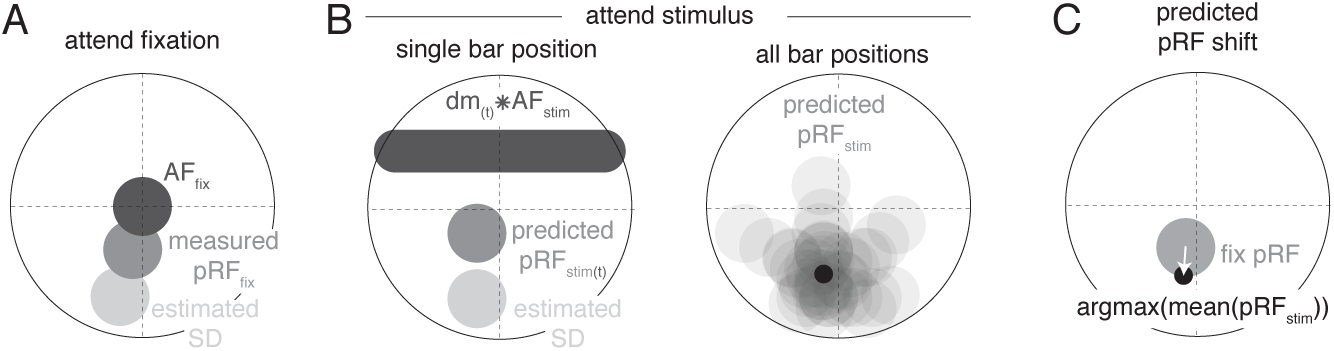
Schematic overview of modeling procedure. A. The Stimulus Drive (SD) was estimated by dividing the measured Attend Fixation pRF by an AF at fixation (*AF_fix_*). B. Attention toward the bar stimulus at a given timepoint *t* was modeled as the multiplication of the estimated SD with the bar stimulus at that timepoint (*dm*_(*t*)_) convolved with another AF (*AF_stim_*). These predicted Attend Stimulus pRFs were averaged over all timepoints. The maximum position of this profile was taken as the predicted Attend Stimulus pRF position. C. The predicted pRF shift ran from the measured Attend Fixation pRF toward the predicted Attend Stimulus position.

**Table 8:** Statistics corresponding to Figure 5D on feature-based attentional modulation. P-values are uncorrected two-tailed tests whether bootstrapped feature attentional modulation index distribution is different from 0. Double and triple asterisks indicate FDR corrected significance of <.01 and <.001 respectively. FDR correction performed over all p-values in this table simultaneously.

| ROI | N | mean diff | p | cohen_d |
| --- | --- | --- | --- | --- |
| V1 | 1605 | 0.05 | <.001*** | 0.170 |
| V2 | 2165 | 0.04 | <.001*** | 0.146 |
| V3 | 1762 | 0.03 | <.001*** | 0.105 |
| hV4 | 860 | 0.05 | <.001*** | 0.197 |
| VO | 451 | 0.08 | <.001*** | 0.290 |
| LO | 1009 | 0.02 | 0.008** | 0.085 |
| V3AB | 703 | 0.11 | <.001*** | 0.364 |
| IPS0 | 237 | 0.14 | <.001*** | 0.486 |
| MT+ | 252 | 0.08 | <.001*** | 0.270 |

**Table 9:**
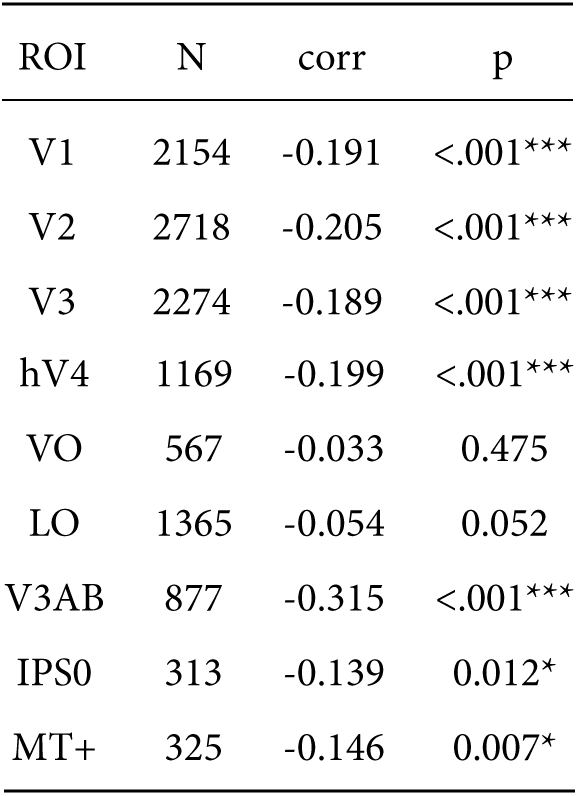
Statistics corresponding to Figure 6 on correlation between feature preference and eccentricity. P-values are uncorrected two-tailed tests whether bootstrapped correlation value of feature preference and pRF eccentricity differs from 0. Single and triple asterisks indicate FDR corrected significance of <.05 and <.001 respectively. FDR correction performed over all p-values in this table simultaneously.

#### Gaze data processing

Gaze data was cleaned by linearly interpolating blinks detected by the EyeLink software. Transient occasions in which the tracker lost the pupil due to partial occlusion by the eyelid leading to high-frequency, high-amplitude signal components were detected and corrected as follows. Pupil size was first high-pass filtered at 10 Hz (the pupil impulse response function is a low-pass filter with a cutoff below 10 Hz (Knapen et al., 2016; Korn and Bach, 2016)), after which those time-points in which the acceleration of pupil size was greater than 10^5^ mm/s, and their neighbours within 5 s, were replaced with NaN values. Drift correction was performed within each bar-pass by subtracting the median gaze position. All gaze positions were rotated to the direction of bar movement, after which we analyzed the median and variance (standard deviation) of the component in the direction of bar movement (i.e. the component relevant for the pRF measurement).

## Acknowledgements

This study was supported in part by an Open Research Area grant (ORA; #464-11-030) issued by the Netherlands Organisation for Scientific Research (NWO) to JT.

## Supplementary Material

### Introduction

This document contains figures and analyses performed for individual subjects. Results are reported using different methods. The ‘super subject’ method pools voxels across subjects and computes p-values and confidence intervals using bootstrapping across voxels. Resulting p-values were FDR corrected for multiple comparisons. This corresponds to what is reported in the main text. The ‘per subject’ method computes these same analyses, but for each subject individually. We report in how many of the subjects these bootstrap tests over voxels are significant with uncorrected p < .05. The ‘over subjects’ method takes the average over the ‘per subject’ results, and computes confidence intervals and t-tests over these 5 values. Statistical tables for all analyses are found at the bottom of this document.

**Figure S1:**
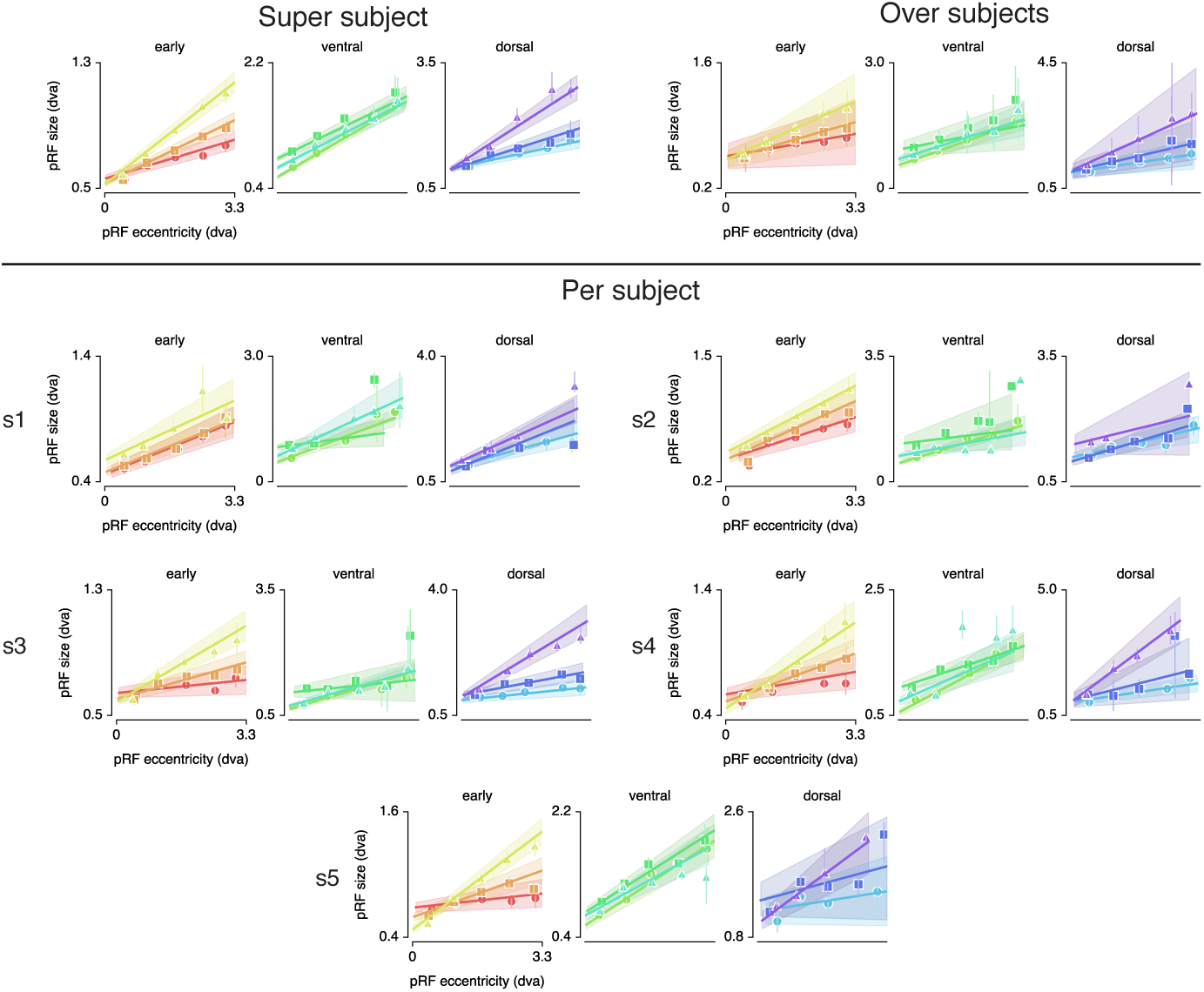
Eccentricity-size relations for all statistical methods.

**Figure S2:**
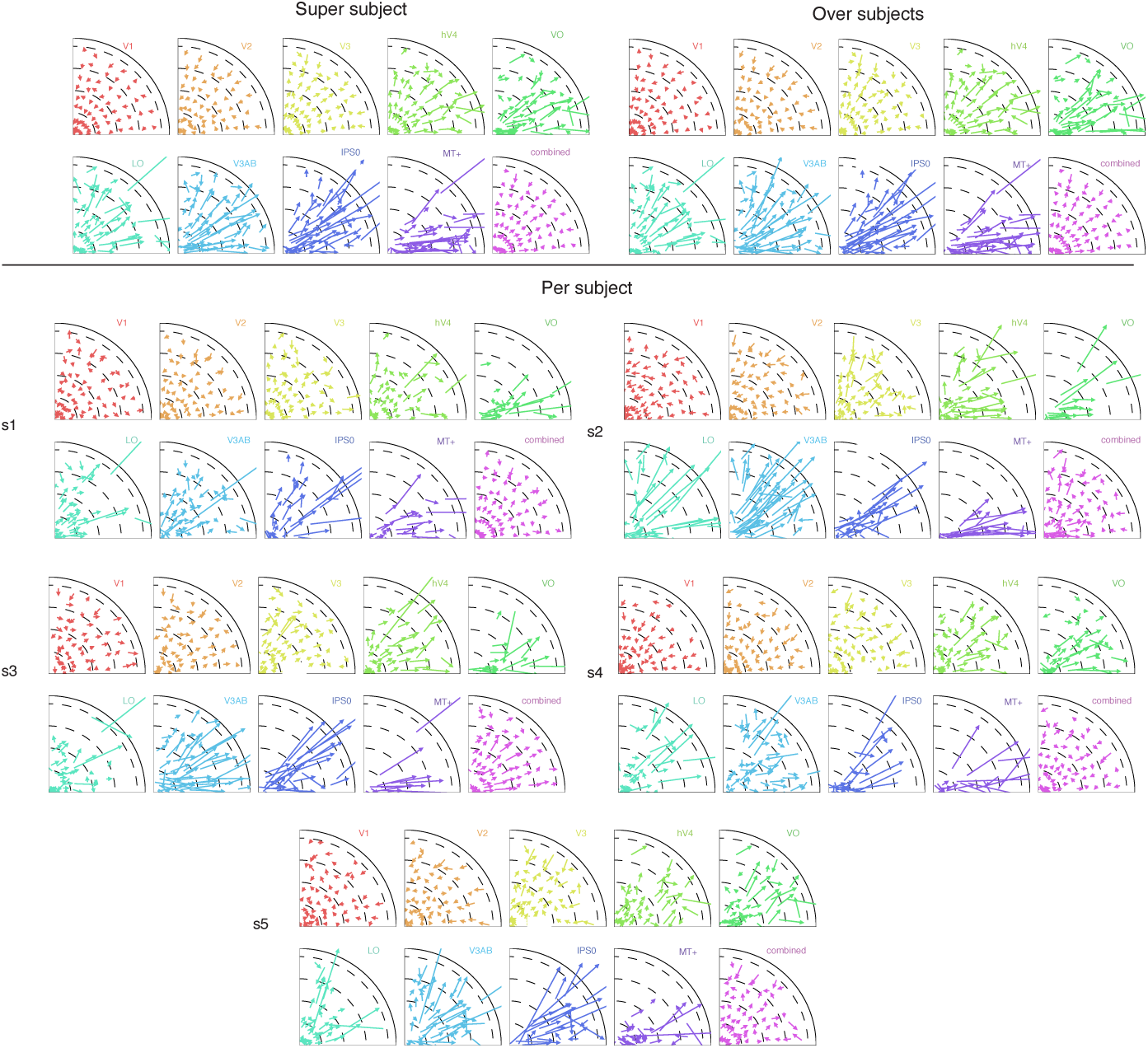
pRF shift plots for the different statistical methods. Shift vectors run from *the Attend Fixation* to *the Attend Stimulus* pRF location. This figure indicates that the shift patterns seen in the super subject are almost identical to the over subjects method, and highly agree with the individual subject figures. Note the radial shift direction that is readily apparant in all ROIs in all subjects. Also note how the *combined* ROI clearly shows paravofeal pRFs shifting toward the periphery and peripheral pRFs shifting toward the fovea across subjects. Finally, the absence of pRFs near the vertical meridian in data from all subjects in VO, IPSO and MT+ highlight the overrepresentation of the horizontal meridian that is found in all ROIs (see Tables 2, 12 and 13.

**Figure S3:**
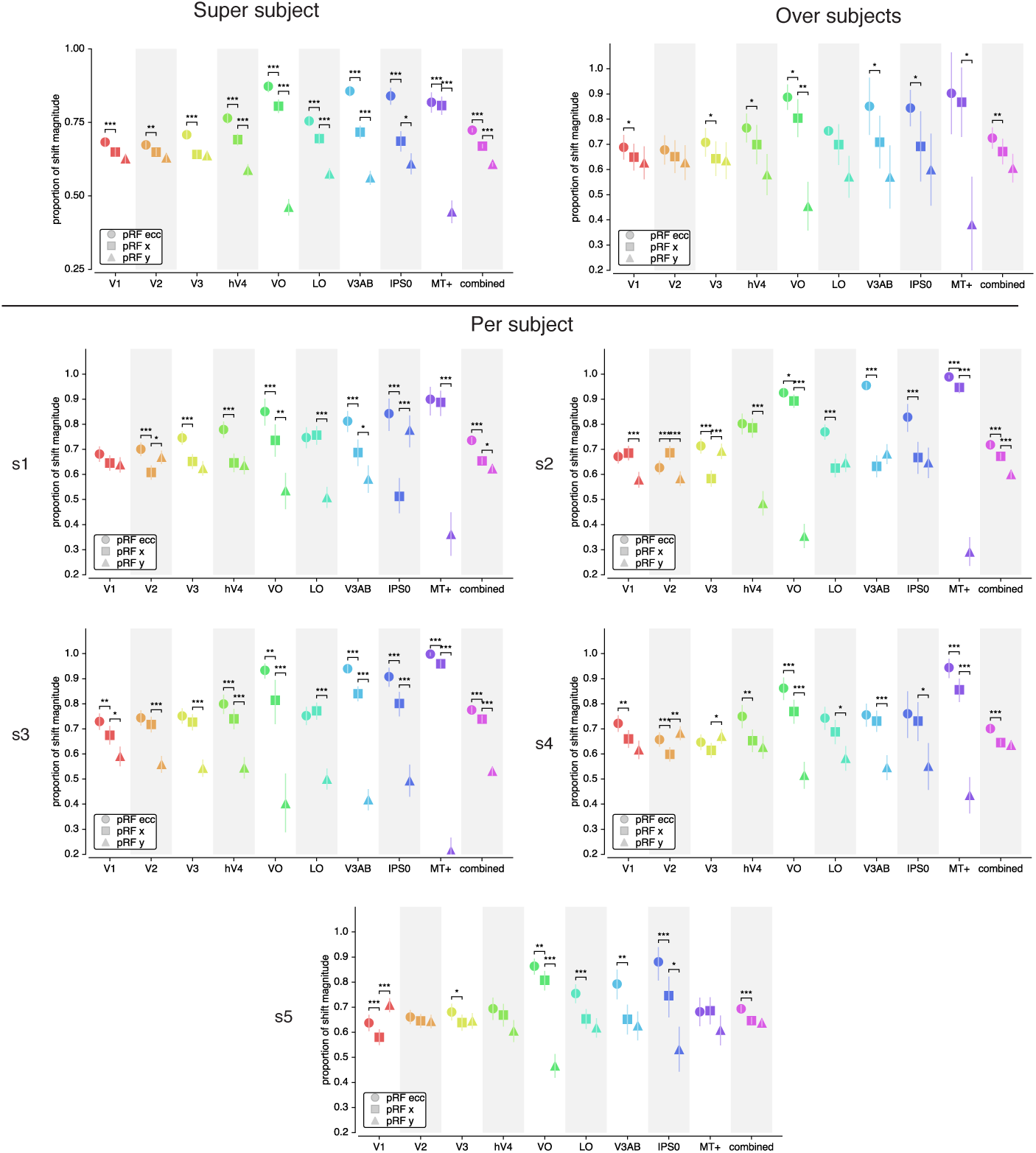
pRF shift directions for the different statistical methods. The ‘over subject’ panel replicates the ‘super subject’ results, namely that shifts are best explained by eccentricity changes, followed by changes in pRF x and finally in pRF y in all ROIs. The dominance of eccentricity over x changes was significant in all individual subjects (’per subject’ method) in the *combined* ROI, and in at least 2 (but often 5) subjects in all other ROIs (see Table 10). The individual subject evidence also replicated the finding that pRFs shifted more in the x compared to y directions in hV4/VO/LO/V3AB/IPSO/MT+ and in the combined ROI (see Table 11). The dominance of x over y shifts was explained from a non-uniformity in the distribution of pRF polar angle (i.e. overrepresentation of horizontal meridian, see Table 2). This was also found in most individual subjects for most ROIs (see Tables 12 and 13). Single, double and tripple asterisks indicate significant differences at p < .05, .01 and .001 respectively.

**Figure S4:**
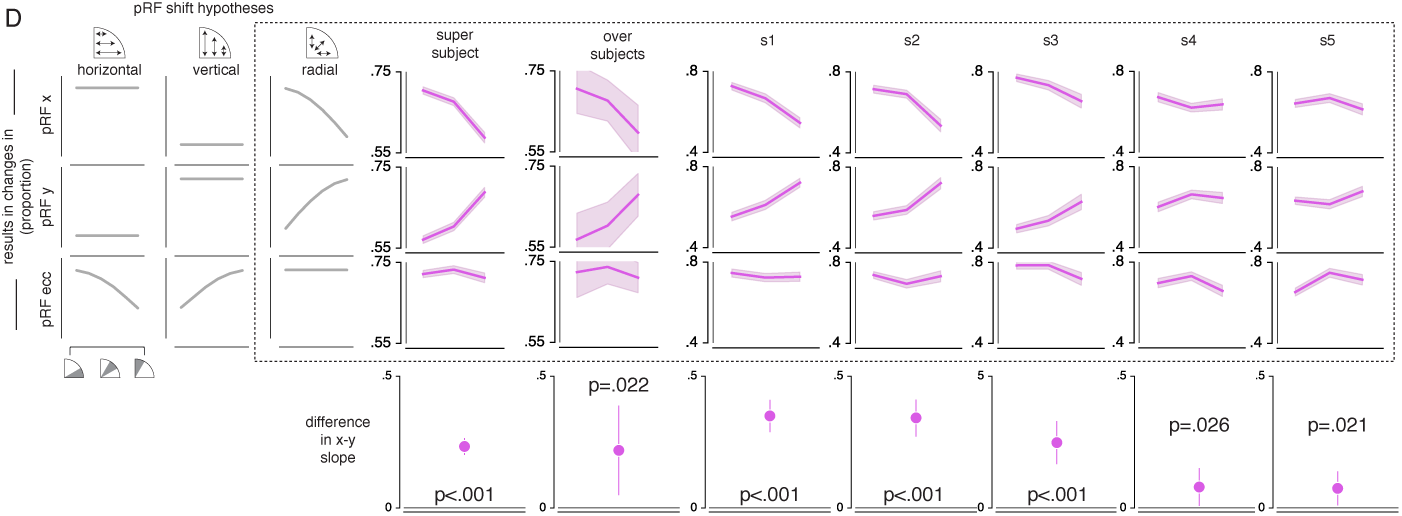
pRF x, y and eccentricity position shifts plotted as a function of polar angle, for different shift direction hypotheses. The data from all statistical methods closely matches the radial shift direction hypothesis, showing strongest pRF x shifts close to the horizontal meridian, strongest pRF y shifts close to the vertical meridian and no polar angle dependence of pRF eccentricity shifts. This is evidenced by a more positive slope of y change over polar angle compared to slope of x change over polar angle in the ‘super subject’ method (see main text for statistics), for the ‘over subjects’ method (slope difference = 0.219, t(4) = 3.616, p = 0.022, cohen’s d = 1.808), and was significant in every individual subject. Error bars reflect 95% CIs across voxels in the ‘super subject’ and ‘per subject’ method, and across subjects in the ‘over subjects’ method.

**Figure S5:**
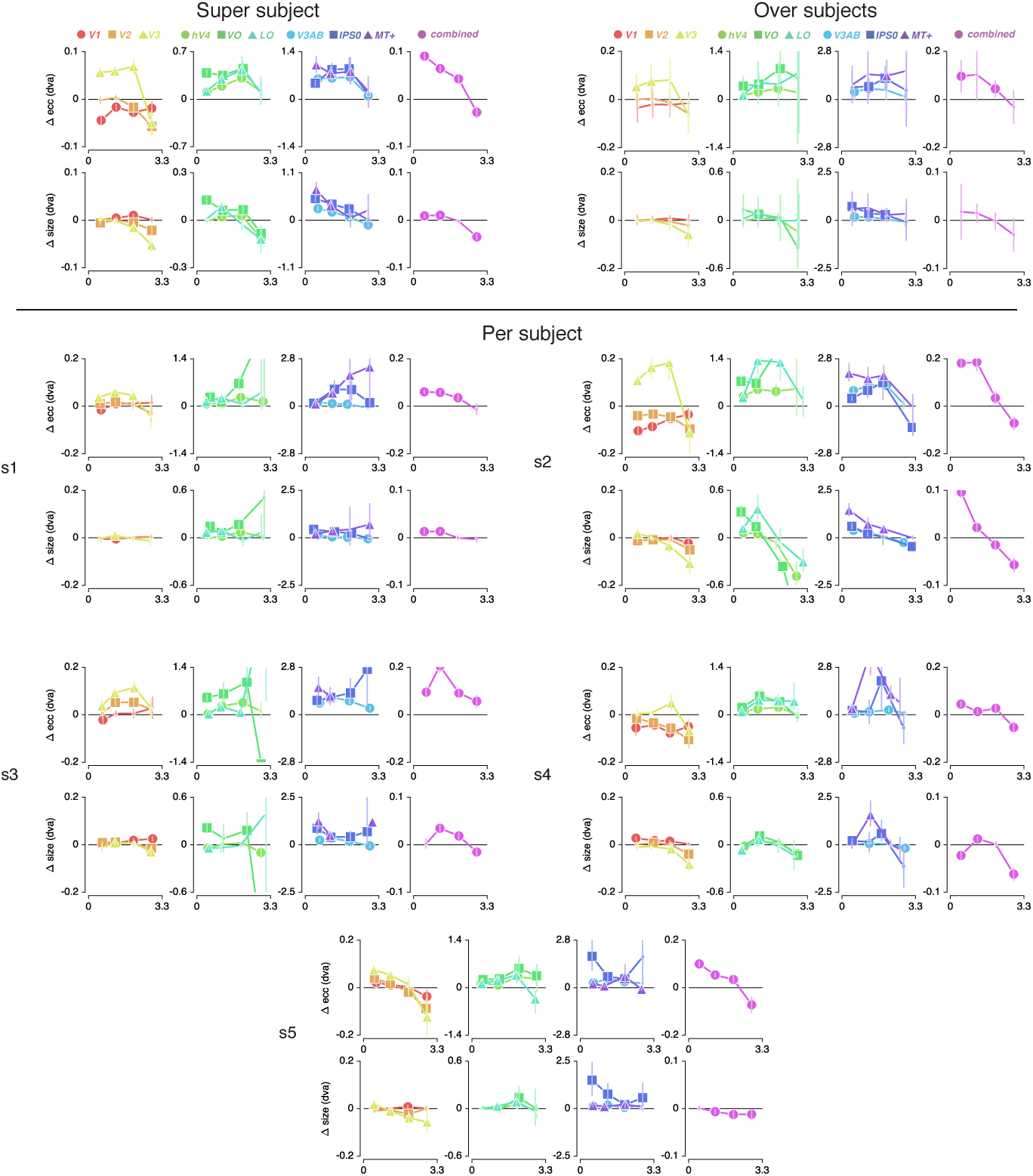
Difference between *Attention to Stimulus* and *Attention to Fixation* pRF eccentricity and size as a function of *Attention to Fixation* eccentricity (see Tables 3–6 and 14–21). The data shows a transition from inwards to outwards shifts across the visual hierarchy in all subjects. Additionally, the progressions of change over eccentricity are similar within ROIs across subjects. Finally, data from the *combined* ROI in most subjects reveals that parafoveal pRFs tend to shift away from the fovea and increase in size, while peripheral pRFs tend to shift toward the fovea and decrease in sizes. In the ‘super subject’ and ‘per subject’ methods markers’ errorbar denotes 95% CI of data over voxels; in the ‘over subjects’ method, errorbar denotes 95% CI over subjects. Increased marker size indicates significance with p < .05 (FDR corrected for ‘super subject’).

**Figure S6:**
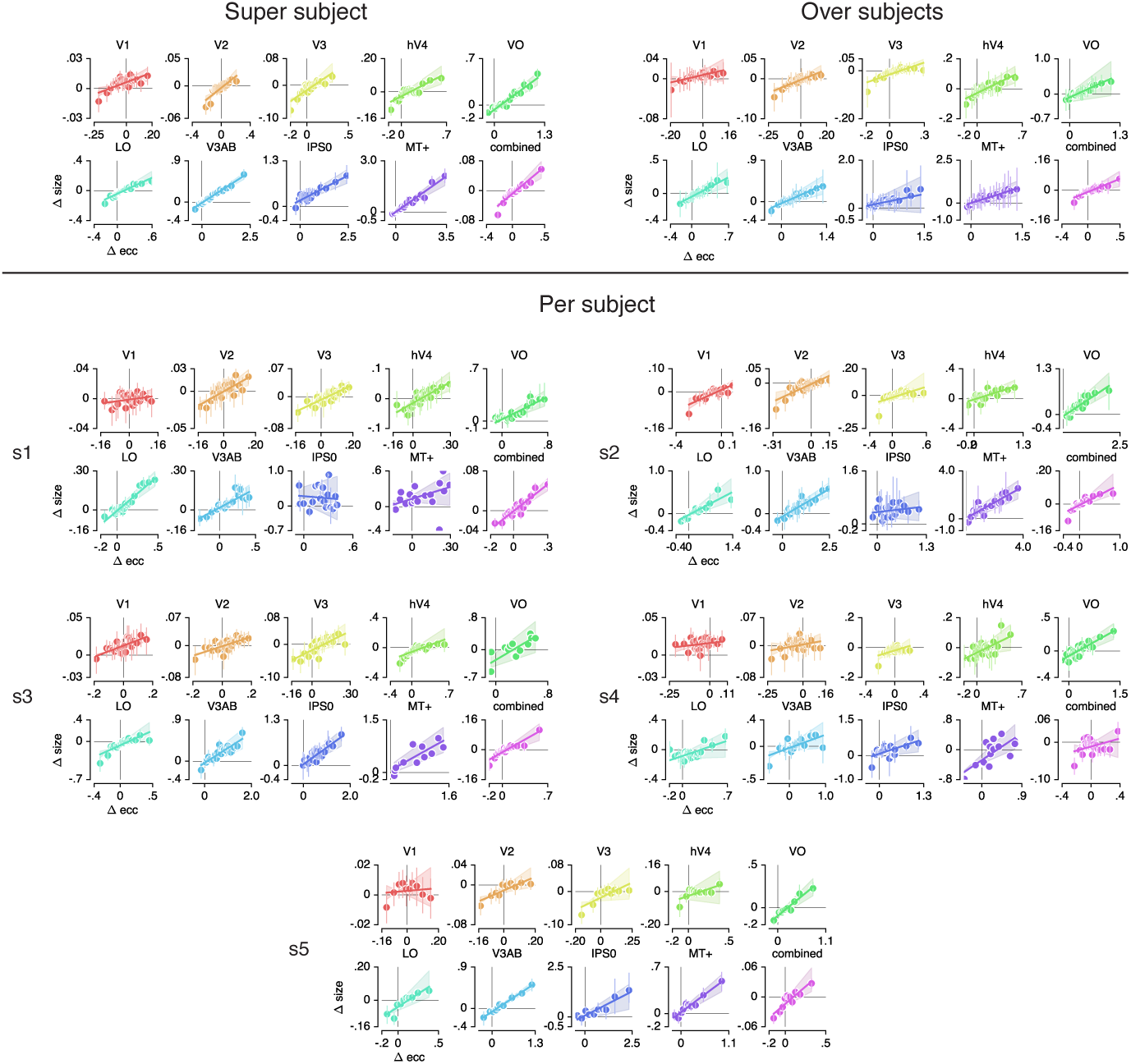
Changes in pRF eccentricity and size were strongly correlated in all ROIs. In the ‘over subjects’ method, we find that pRF eccentricity and size changes are significant in all ROIs except IPSO (although p = .070). In the ‘per subject method’, we find such a significant correlation in at least 2 (but often 5) subjects (see Table 22). In the ‘super subject’ and for ‘per subjects’ figures, markers’ errorbar denotes 95% CI of data over voxels; in the ‘over subjects’ figure, errorbar denotes 95% CI over subjects.

**Figure S7:**
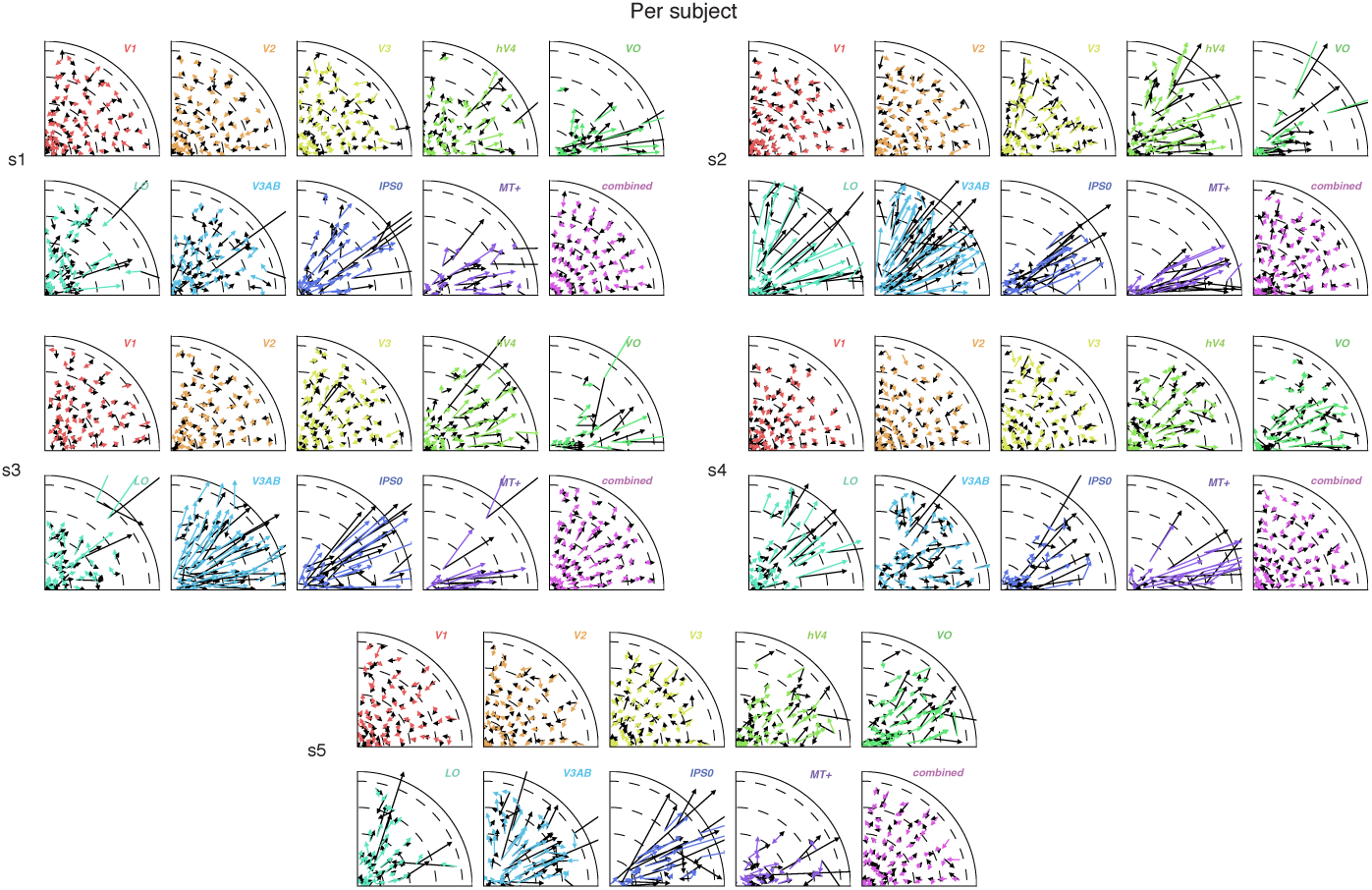
Attentional gain field modeling results for each subject. Arrows depict observed (black) and predicted (color) pRF shifts. This shows that the model closely captured the data in all individual subjects.

**Figure S8:**
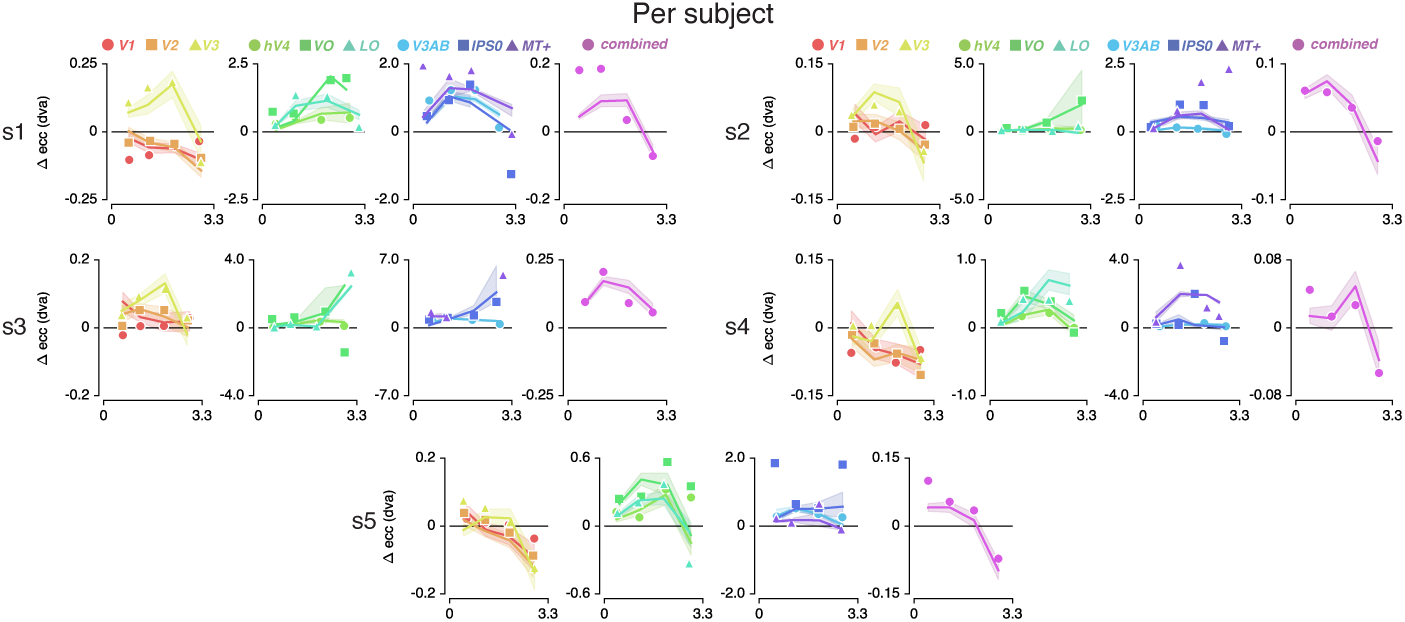
Attentional gain field modeling results for each subject. Observed (markers) and predicted (solid lines) pRF eccentricity difference between *Attend Fixation* and *Attend Stimulus* conditions for each subject. Shaded areas indicate 95% CI over voxels. This shows that the model was able to capture between subject variance with a high accuracy.

**Figure S9:**
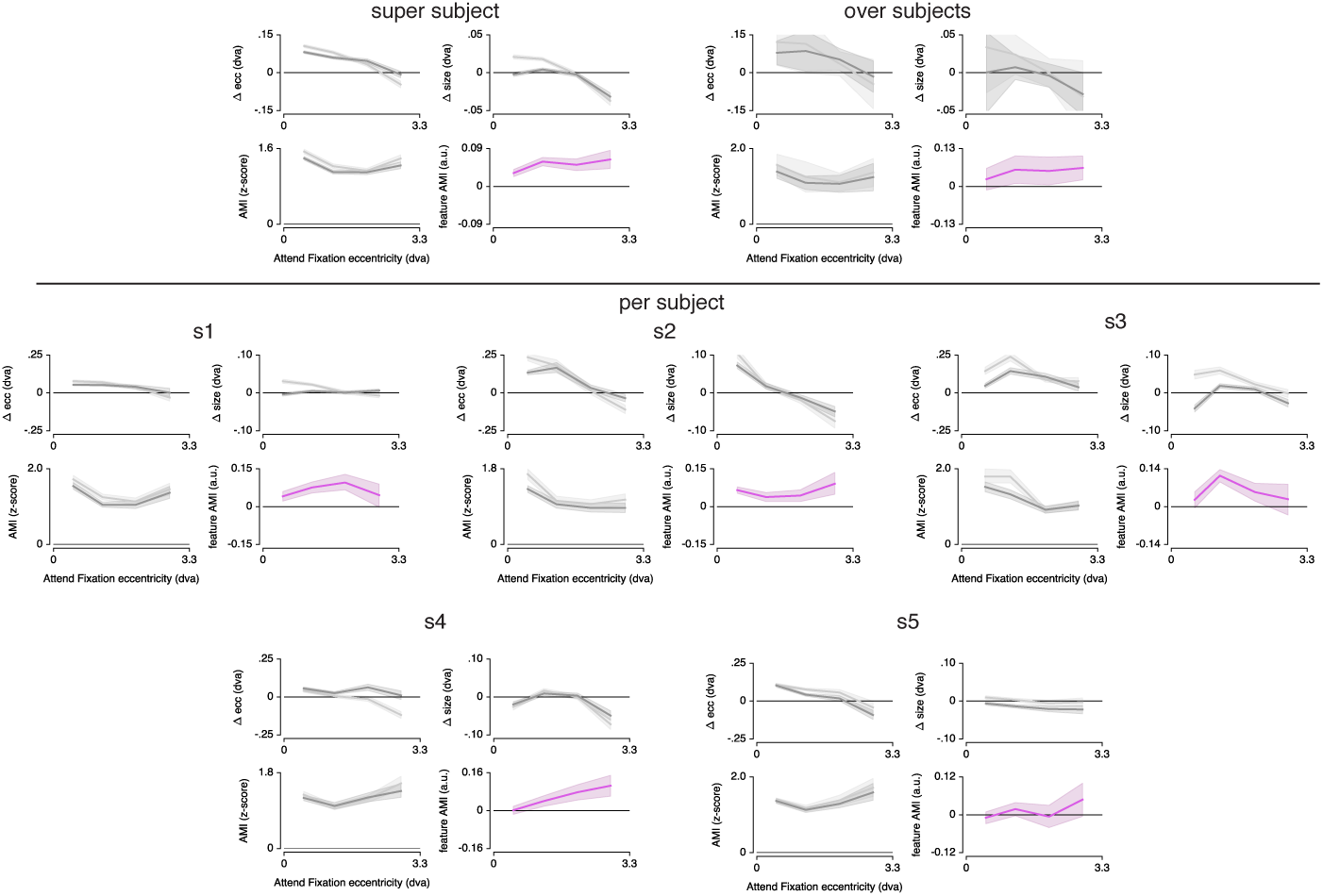
Differences in pRF eccentricity and size relative to the *Attention to Fixation* condition, for both *the Attention to Color* and *Attention to TF* condition separately. The changes in both eccentricity and size are more pronounced when attending changes in color versus TF in the bar. This pattern is most clear in the ‘super subject’ and ‘over subjects’ method (albeit with larger variance), and is present in most individual subjects. In the ‘super subject’ and ‘per subject’ figures, error bars denote 95% CI over voxels, in the ‘over subject’ method error bars indicate 95% CI over subjects.

**Figure S10:**
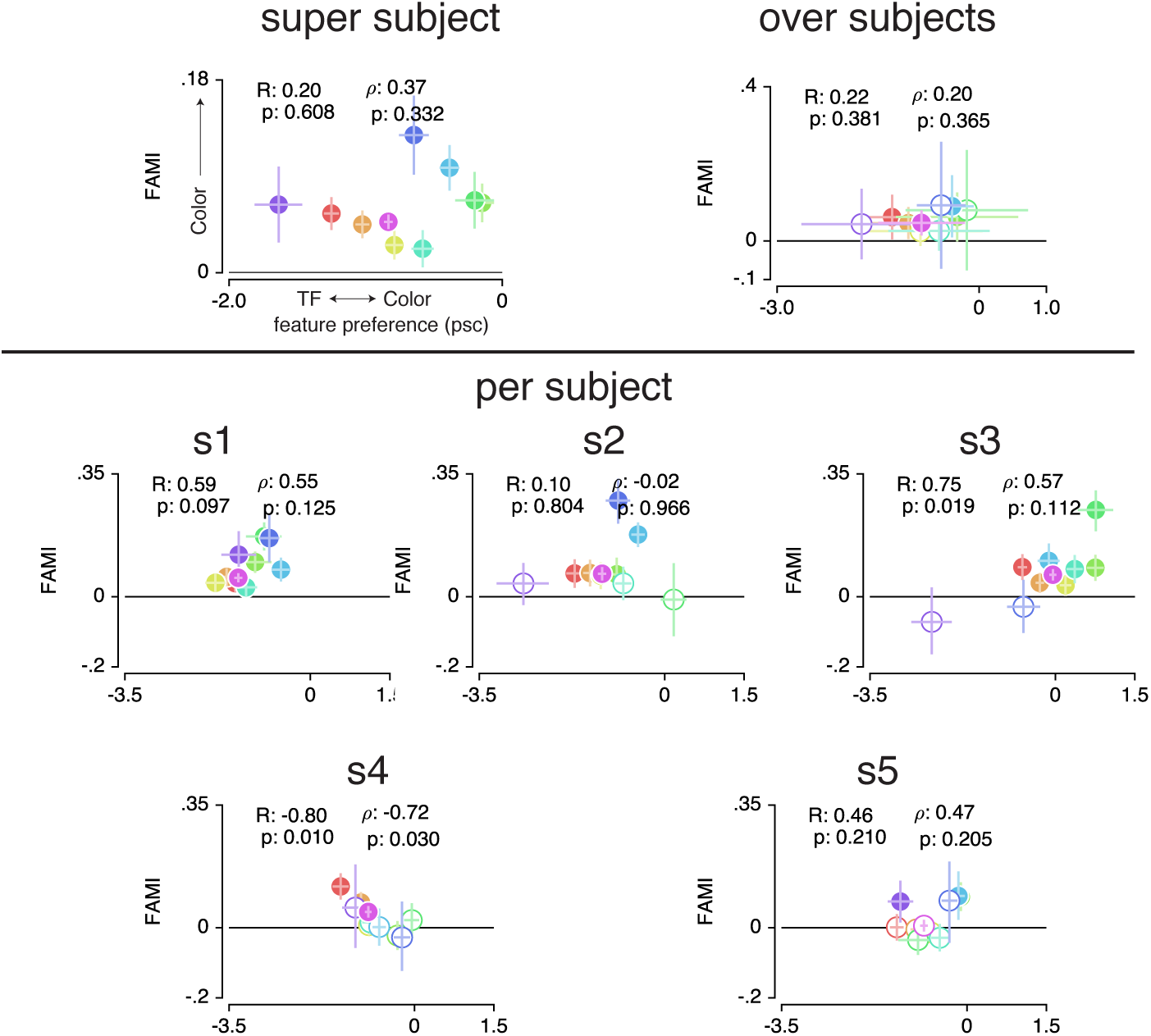
Feature AMI compared to feature preference for each ROI, for each statistical method. The y-axis displays feature AMI, which increases when pRF modulations (size and eccentricity changes combined, see Methods) are greater when attending color compared to TF. The x-axis displays feature preference, which inreases with higher color compared to TF preference. pRF modulations were greater when attending color in all ROIs in the ‘super subject’ method, and was unrelated to feature preference (see Table 8). In the ‘over subjects’ method, we confirm that feature AMI was not different between ROIs over subjects (RM ANOVA, factor of ROI: F(8,4) = 1.108, p = 0.384, *η^2^ρ =* 0.225), and that it was on average 0.064 over ROIs across subjects, t(4) = 4.654, p = 0.010, cohen’s d = 2.081). Additionally, we found that feature AMI was significantly positive in at least 2 (but often 4) out of 5 subjects in each ROI (see Table 23). Finally, we also found no correlation with feature preference in the ‘over subjects’ method (t(4) = 0.655, p = .549, cohen’s d = 0.33).

**Figure S11:**
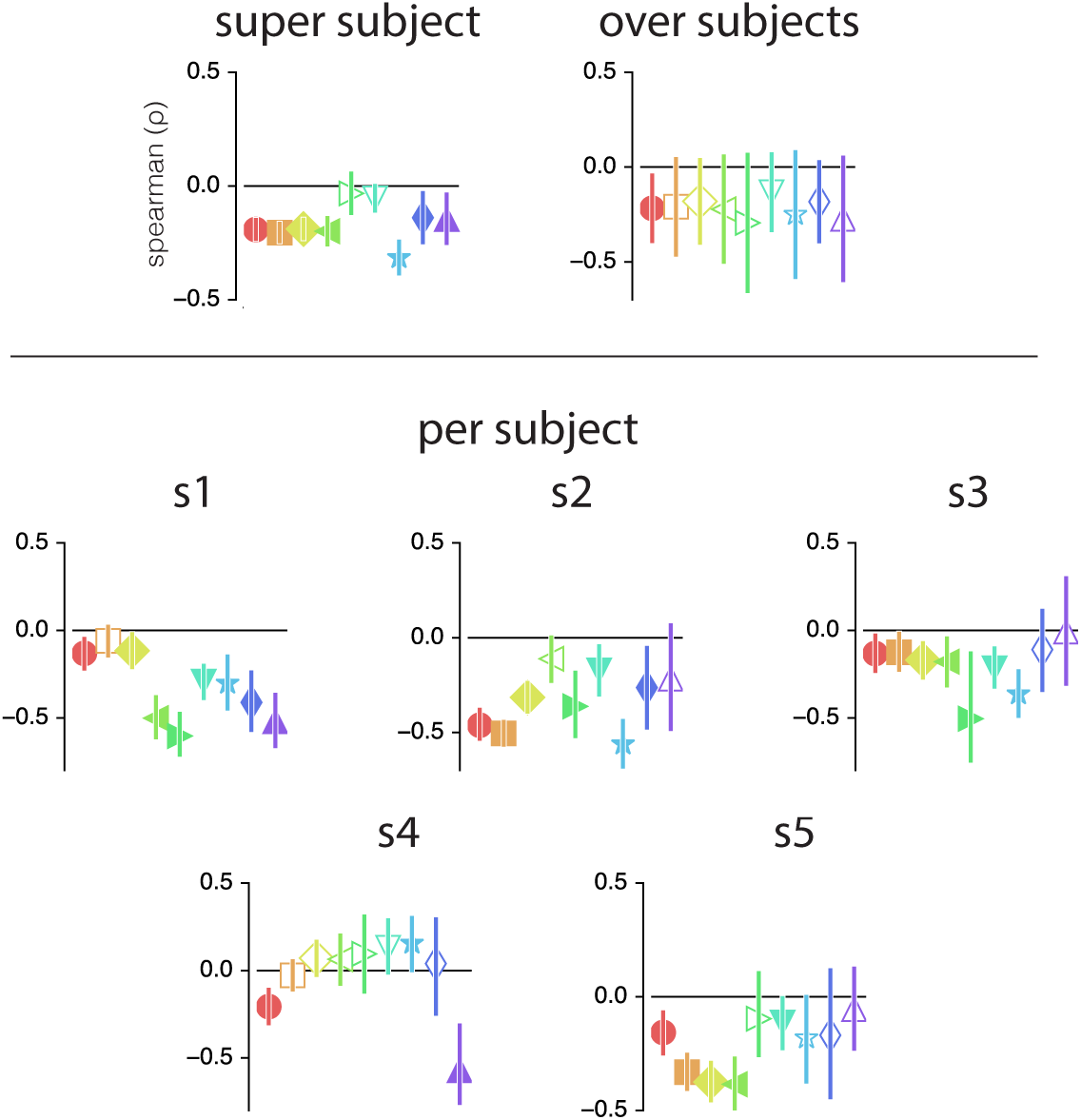
Color compared to TF preference versus eccentricity correlations. The ‘super subject’ method shows negative correlations in all ROIs except VO and LO. The ‘over’ subjects method shows that this correlation is negative on average and does not vary across ROI (RM ANOVA with main factor of ROI F(8,4) = 0.354, p = .937, *η*^2^*ρ = .030*, on average −0.218 over ROIs, F(1,4) = 15.630, p = .017, *η*^2^*ρ =* .641). When looking at individual subjects, we find that correlations are positive in at least two but often in the majority of subjects across ROIs (see Table 24).

### Statistical Tables - Supplementary Material

**Table 10:** Statistics corresponding to Supplementary Figure 3 on pRF x versus y shift ratios. P-values result from t-tests over the 5 subject values. Single, double and triple asterisks indicates uncorrected p-value < .05, .01 and .001 respectively. The last two columns reflect in how many out of 5 subjects these differences were different from 0 with uncorrected bootstrapped p-values of <.05 over voxels.

| ROI | N | p | t | $\Delta$ ratio | cohen's d | x>y | y<x |
| --- | --- | --- | --- | --- | --- | --- | --- |
| V1 | 5 | .614 | 0.546 | .02 | .27 | 2 | 1 |
| V2 | 5 | .638 | 0.508 | .02 | .25 | 2 | 2 |
| V3 | 5 | .882 | 0.158 | .01 | .08 | 1 | 2 |
| hV4 | 5 | .100 | 2.128 | .12 | 1.06 | 2 | 0 |
| VO | 5 | .004** | 5.858 | .35 | 2.93 | 5 | 0 |
| LO | 5 | .091 | 2.216 | .13 | 1.11 | 3 | 0 |
| V3AB | 5 | .165 | 1.699 | .14 | .085 | 3 | 0 |
| IPS0 | 5 | .411 | 0.918 | .09 | .46 | 3 | 1 |
| MT+ | 5 | .014* | 4.162 | .48 | 2.08 | 4 | 0 |
| combined | 5 | .151 | 1.773 | .07 | .89 | 3 | 0 |

**Table 11:** Statistics corresponding to Supplementary Figure 3 on pRF eccentricity versus x shift ratios. P-values result from t-tests over the 5 subject values. Single, double and triple asterisks indicates uncorrected p-value < .05, .01 and .001 respectively. The last two columns reflect in how many out of 5 subjects these differences were different from 0 with uncorrected bootstrapped p-values of <.05 over voxels.

| ROI | N | p | t | $\Delta$ ratio | cohen's d | ecc>x | ecc<x |
| --- | --- | --- | --- | --- | --- | --- | --- |
| V1 | 5 | .049* | 2.797 | .04 | 1.40 | 3 | 0 |
| V2 | 5 | .350 | 1.057 | .03 | .53 | 2 | 1 |
| V3 | 5 | .036* | 3.094 | .06 | 1.55 | 3 | 0 |
| hV4 | 5 | .037* | 3.072 | .07 | 1.54 | 3 | 0 |
| VO | 5 | .013* | 4.240 | .08 | 2.12 | 5 | 0 |
| LO | 5 | .153 | 1.764 | .06 | .88 | 2 | 0 |
| V3AB | 5 | .037* | 3.084 | .14 | 1.54 | 4 | 0 |
| IPS0 | 5 | .047* | 2.845 | .15 | 1.42 | 4 | 0 |
| MT+ | 5 | .080 | 2.335 | .03 | 1.17 | 3 | 0 |
| combined | 5 | .003** | 6.707 | .05 | 3.35 | 5 | 0 |

**Table 12:** Rayleigh test for non-uniformity for subjects 1-3. Single, double and triple asterisks indicates uncorrected p-value <.05, .01 and .001 respectively.

|  | s1 |  |  | s2 |  |  | s3 |  |  |
| --- | --- | --- | --- | --- | --- | --- | --- | --- | --- |
| ROI | N | z | p | N | z | p | N | z | p |
| V1 | 469 | 3.221 | .040* | 558 | 26.131 | <.001*** | 333 | 11.823 | <.001*** |
| V2 | 592 | 0.659 | .518 | 605 | 9.986 | <.001*** | 411 | 48.090 | <.001*** |
| V3 | 533 | 1.021 | .360 | 463 | 22.909 | <.001*** | 389 | 12.275 | <.001*** |
| hV4 | 335 | 18.664 | <.001*** | 183 | 0.626 | .535 | 228 | 32.801 | <.001*** |
| VO | 97 | 37.899 | <.001*** | 136 | 46.282 | <.001*** | 36 | 11.037 | <.001*** |
| LO | 271 | 2.255 | .105 | 367 | 25.638 | <.001*** | 268 | 37.663 | <.001*** |
| V3AB | 169 | 4.560 | .010* | 176 | 23.160 | <.001*** | 233 | 43.613 | <.001*** |
| IPS0 | 67 | 0.101 | .905 | 77 | 17.517 | <.001*** | 78 | 38.176 | <.001*** |
| MT+ | 43 | 14.066 | <.001*** | 71 | 17.095 | <.001*** | 43 | 27.463 | <.001*** |
| combined | 2576 | 12.585 | <.001*** | 2636 | 119.207 | <.001*** | 2019 | 208.441 | <.001*** |

**Table 13:** Rayleigh test for non-uniformity for subjects 4-5. Single, double and triple asterisks indicates uncorrected p-value <.05, .01 and .001 respectively.

|  | s4 |  |  | s5 |  |  |
| --- | --- | --- | --- | --- | --- | --- |
| ROI | N | z | p | N | z | p |
| V1 | 369 | 3.891 | .020* | 447 | 40.575 | <.001*** |
| V2 | 563 | 4.195 | .015* | 583 | 20.792 | <.001*** |
| V3 | 504 | 22.465 | <.001*** | 433 | 27.032 | <.001*** |
| hV4 | 218 | 0.995 | .370 | 237 | 30.059 | <.001*** |
| VO | 146 | 24.882 | <.001*** | 167 | 28.314 | <.001*** |
| LO | 195 | 4.618 | .010* | 304 | 24.695 | <.001*** |
| V3AB | 176 | 3.471 | .031* | 129 | 18.098 | <.001*** |
| IPS0 | 49 | 12.339 | <.001*** | 45 | 11.335 | <.001*** |
| MT+ | 47 | 26.131 | <.001*** | 124 | 50.139 | <.001*** |
| combined | 2267 | 51.292 | <.001*** | 2469 | 163.225 | <.001*** |

**Table 14:** Statistics corresponding to Supplementary Figure 5 on pRF eccentricity changes in eccentricity bin 1. P-values reflect whether t-test showed that subject average values were different from 0, for each ROI. Single, double and triple asterisks indicate uncorrected significance of <.05, <.01 and <.001 respectively. Right most columns indicate in how many out of 5 subjects the bootstrap test over voxels was different from 0 with uncorrect p-value of < .05.

| ROI | N | p | t | $\Delta$ ecc | cohen's d | pos in | neg in |
| --- | --- | --- | --- | --- | --- | --- | --- |
| V1 | 5 | .172 | -1.663 | -0.035 | -0.831 | 1 | 4 |
| V2 | 5 | .998 | 0.003 | 0.000 | 0.001 | 2 | 2 |
| V3 | 5 | .042* | 2.953 | 0.053 | 1.477 | 4 | 0 |
| hV4 | 5 | .052 | 2.729 | 0.124 | 1.365 | 5 | 0 |
| VO | 5 | .017* | 3.903 | 0.393 | 1.952 | 5 | 0 |
| LO | 5 | .030* | 3.309 | 0.125 | 1.655 | 5 | 0 |
| V3AB | 5 | .047* | 2.843 | 0.442 | 1.422 | 5 | 0 |
| IPS0 | 5 | .080 | 2.335 | 0.715 | 1.167 | 5 | 0 |
| MT+ | 5 | .089 | 2.237 | 0.858 | 1.119 | 5 | 0 |
| combined | 5 | .015* | 4.067 | 0.096 | 2.033 | 5 | 0 |

**Table 15:**
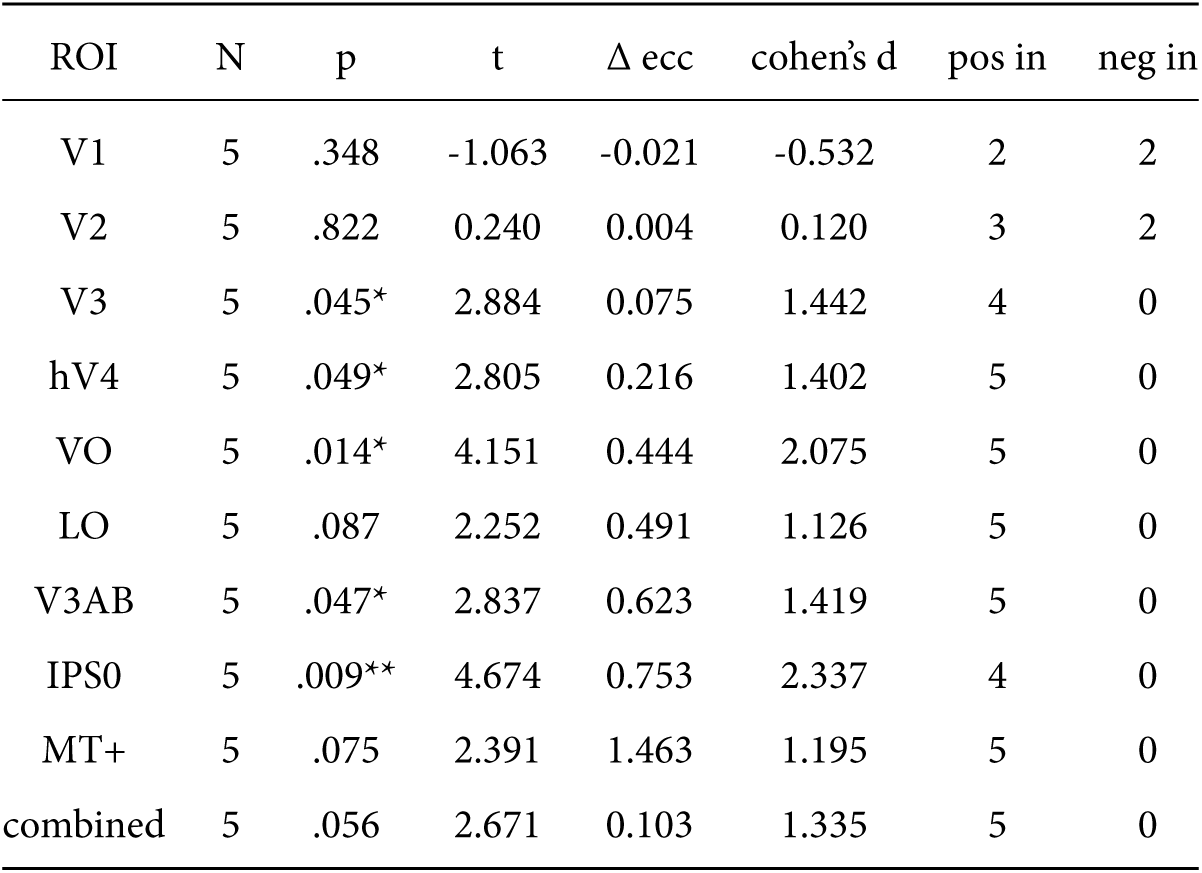
Statistics corresponding to Supplementary Figure 5 on pRF eccentricity changes in eccentricity bin 2. P-values reflect whether t-test showed that subject average values were different from 0, for each ROI. Single, double and triple asterisks indicate uncorrected significance of <.05, <.01 and <.001 respectively. Right most columns indicate in how many out of 5 subjects the bootstrap test over voxels was different from 0 with uncorrect p-value of < .05.

**Table 16:**
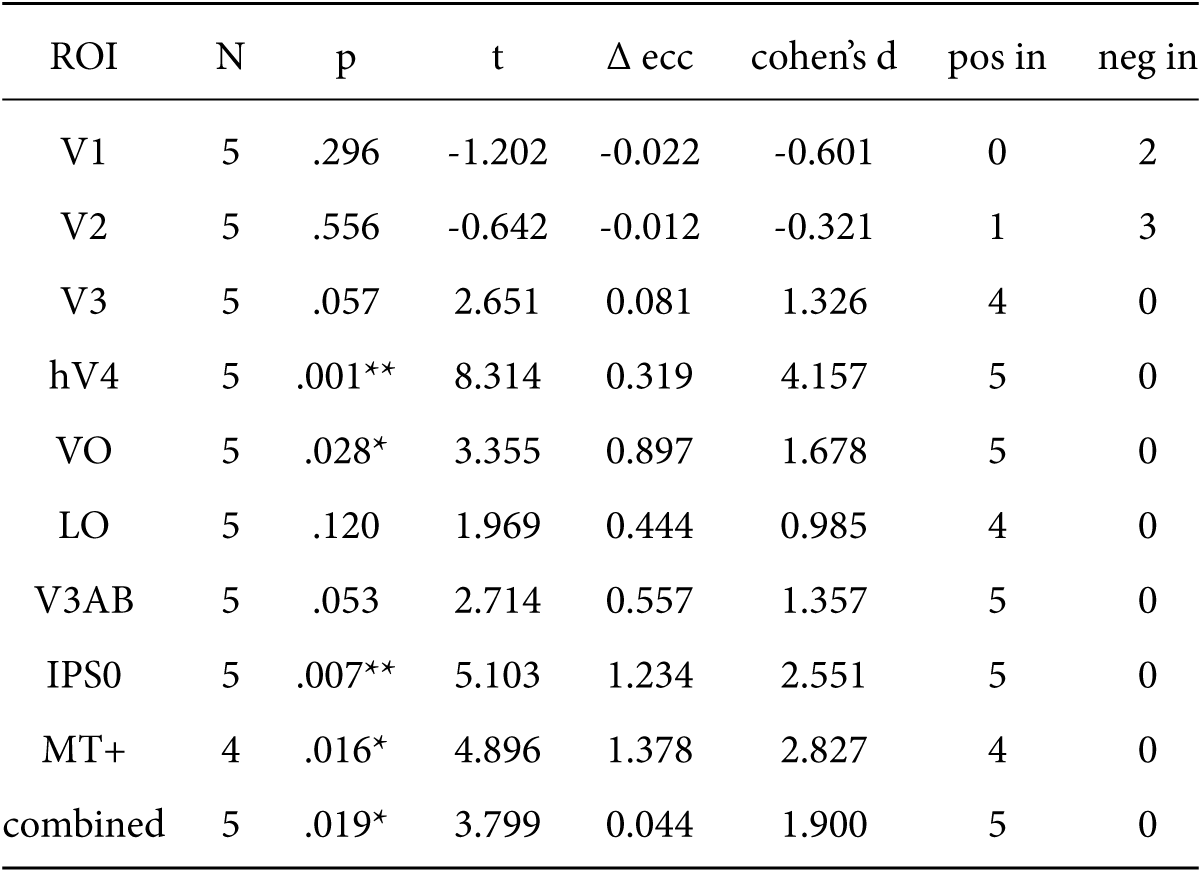
Statistics corresponding to Supplementary Figure 5 on pRF eccentricity changes in eccentricity bin 3. P-values reflect whether t-test showed that subject average values were different from 0, for each ROI. Single, double and triple asterisks indicate uncorrected significance of <.05, <.01 and <.001 respectively. Right most columns indicate in how many out of 5 subjects the bootstrap test over voxels was different from 0 with uncorrect p-value of < .05.

| ROI | N | p | t | $\Delta$ ecc | cohen's d | pos in | neg in |
| --- | --- | --- | --- | --- | --- | --- | --- |
| V1 | 5 | .296 | -1.202 | -0.022 | -0.601 | 0 | 2 |
| V2 | 5 | .556 | -0.642 | -0.012 | -0.321 | 1 | 3 |
| V3 | 5 | .057 | 2.651 | 0.081 | 1.326 | 4 | 0 |
| hV4 | 5 | .001** | 8.314 | 0.319 | 4.157 | 5 | 0 |
| VO | 5 | .028* | 3.355 | 0.897 | 1.678 | 5 | 0 |
| LO | 5 | .120 | 1.969 | 0.444 | 0.985 | 4 | 0 |
| V3AB | 5 | .053 | 2.714 | 0.557 | 1.357 | 5 | 0 |
| IPS0 | 5 | .007** | 5.103 | 1.234 | 2.551 | 5 | 0 |
| MT+ | 4 | .016* | 4.896 | 1.378 | 2.827 | 4 | 0 |
| combined | 5 | .019* | 3.799 | 0.044 | 1.900 | 5 | 0 |

**Table 17:** Statistics corresponding to Supplementary Figure 5 on pRF eccentricity changes in eccentricity bin 4. P-values reflect whether t-test showed that subject average values were different from 0, for each ROI. Single, double and triple asterisks indicate uncorrected significance of <.05, <.01 and <.001 respectively. Right most columns indicate in how many out of 5 subjects the bootstrap test over voxels was different from 0 with uncorrect p-value of < .05.

| ROI | N | p | t | $\Delta$ ecc | cohen's d | pos in | neg in |
| --- | --- | --- | --- | --- | --- | --- | --- |
| V1 | 5 | .369 | -1.011 | -0.016 | -0.506 | 0 | 3 |
| V2 | 5 | .071 | -2.442 | -0.059 | -1.221 | 0 | 3 |
| V3 | 5 | .085 | -2.273 | -0.063 | -1.136 | 0 | 3 |
| hV4 | 5 | .082 | 2.308 | 0.202 | 1.154 | 1 | 0 |
| VO | 5 | .417 | 0.904 | 0.623 | 0.452 | 3 | 1 |
| LO | 5 | .285 | 1.234 | 0.782 | 0.617 | 2 | 1 |
| V3AB | 5 | .122 | 1.960 | 0.154 | 0.980 | 1 | 0 |
| IPS0 | 5 | .516 | 0.711 | 0.533 | 0.356 | 2 | 1 |
| MT+ | 4 | .187 | 1.590 | 1.662 | 0.795 | 2 | 1 |
| combined | 5 | .273 | -1.270 | -0.031 | -0.635 | 1 | 3 |

**Table 18:**
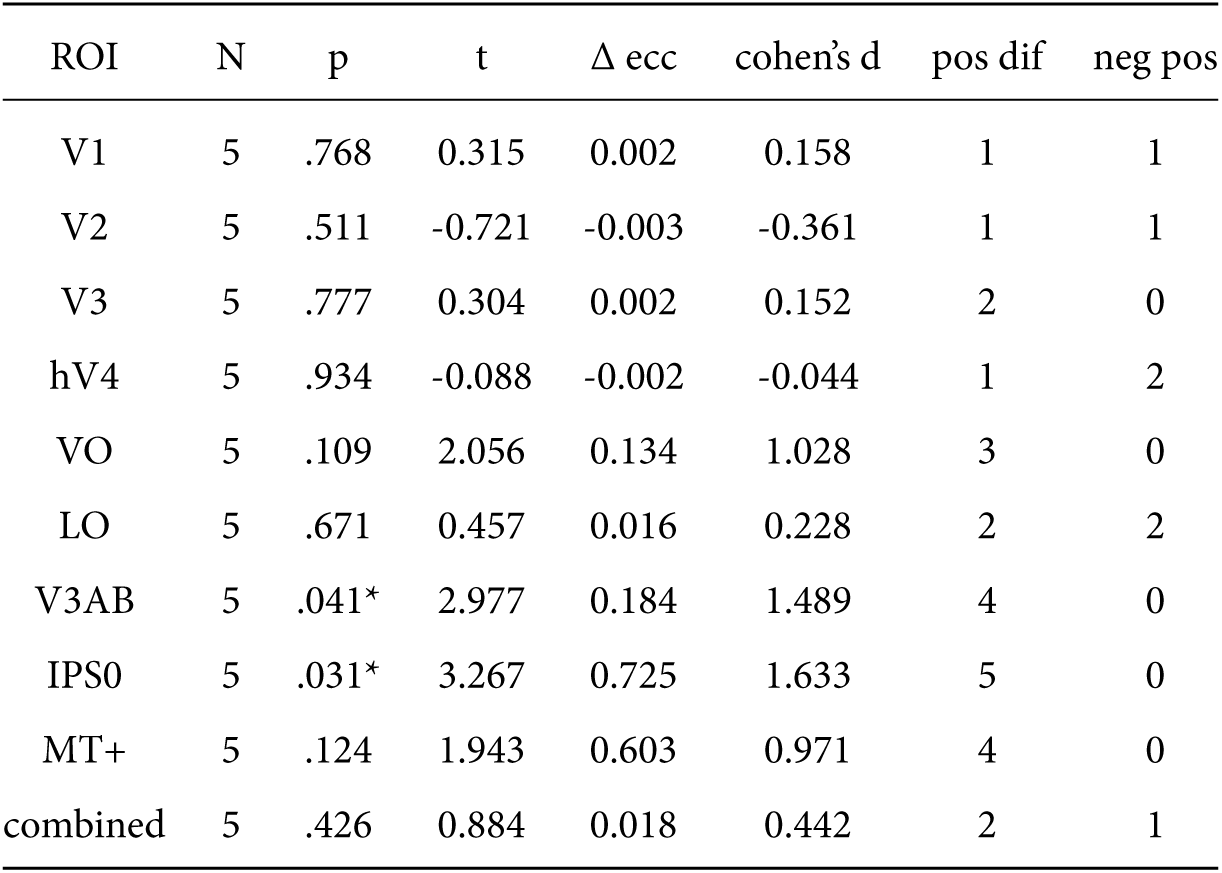
Statistics corresponding to Supplementary Figure 5 on pRF size changes in eccentricity bin 1. P-values reflect whether t-test showed that subject average values were different from 0, for each ROI. Single, double and triple asterisks indicate uncorrected significance of <.05, <.01 and <.001 respectively. Right most columns indicate in how many out of 5 subjects the bootstrap test over voxels was different from 0 with uncorrect p-value of < .05.

**Table 19:** Statistics corresponding to Supplementary Figure 5 on pRF size changes in eccentricity bin 2. P-values reflect whether t-test showed that subject average values were different from 0, for each ROI. Single, double and triple asterisks indicate uncorrected significance of <.05, <.01 and <.001 respectively. Right most columns indicate in how many out of 5 subjects the bootstrap test over voxels was different from 0 with uncorrect p-value of < .05.

| ROI | N | p | t | $\Delta$ ecc | cohen's d | pos dif | neg pos |
| --- | --- | --- | --- | --- | --- | --- | --- |
| V1 | 5 | .572 | 0.614 | 0.003 | 0.307 | 2 | 1 |
| V2 | 5 | .697 | 0.418 | 0.002 | 0.209 | 2 | 1 |
| V3 | 5 | .937 | 0.085 | 0.000 | 0.042 | 2 | 1 |
| hV4 | 5 | .082 | 2.308 | 0.024 | 1.154 | 3 | 0 |
| VO | 5 | .037* | 3.089 | 0.078 | 1.544 | 3 | 0 |
| LO | 5 | .214 | 1.475 | 0.101 | 0.738 | 4 | 0 |
| V3AB | 5 | .033* | 3.211 | 0.175 | 1.605 | 5 | 0 |
| IPS0 | 5 | .027* | 3.419 | 0.367 | 1.709 | 4 | 0 |
| MT+ | 5 | .058 | 2.631 | 0.656 | 1.316 | 5 | 0 |
| combined | 5 | .088 | 2.250 | 0.015 | 1.125 | 4 | 1 |

**Table 20:** Statistics corresponding to Supplementary Figure 5 on pRF size changes in eccentricity bin 3. P-values reflect whether t-test showed that subject average values were different from 0, for each ROI. Single, double and triple asterisks indicate uncorrected significance of <005, <.01 and <.001 respectively. Right most columns indicate in how many out of 5 subjects the bootstrap test over voxels was different from 0 with uncorrect p-value of < .05.

| ROI | N | p | t | $\Delta$ ecc | cohen's d | pos dif | neg pos |
| --- | --- | --- | --- | --- | --- | --- | --- |
| V1 | 5 | .061 | 2.579 | 0.009 | 1.290 | 3 | 0 |
| V2 | 5 | .373 | -1.001 | -0.005 | -0.501 | 0 | 1 |
| V3 | 5 | .111 | -2.041 | -0.017 | -1.020 | 0 | 3 |
| hV4 | 5 | .376 | 0.995 | 0.022 | 0.498 | 1 | 0 |
| VO | 5 | .773 | 0.308 | 0.031 | 0.154 | 3 | 1 |
| LO | 5 | .800 | -0.271 | -0.007 | -0.135 | 1 | 0 |
| V3AB | 5 | .085 | 2.280 | 0.068 | 1.140 | 3 | 0 |
| IPS0 | 5 | .034* | 3.164 | 0.297 | 1.582 | 4 | 0 |
| MT+ | 4 | .076 | 2.662 | 0.285 | 1.537 | 2 | 0 |
| combined | 5 | .800 | -0.270 | -0.002 | -0.135 | 1 | 2 |

**Table 21:**
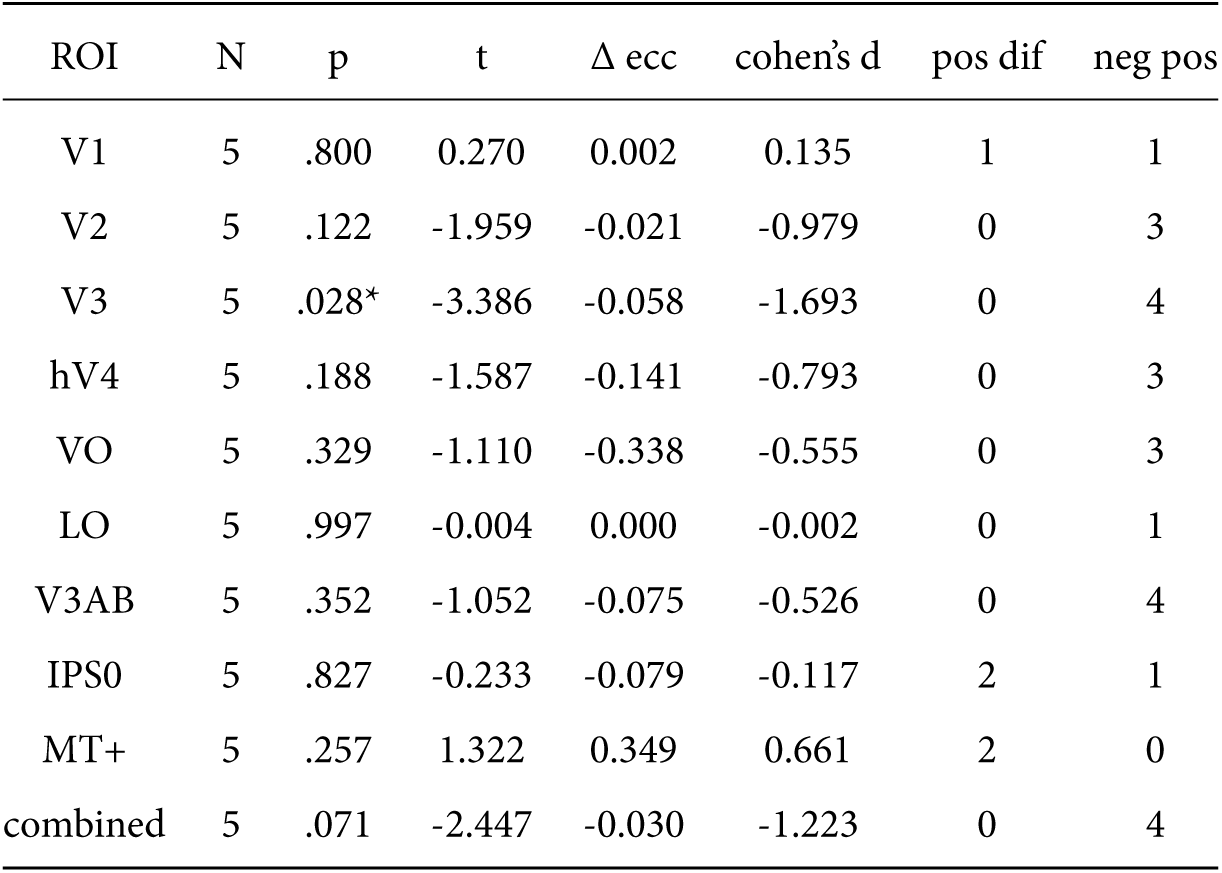
Statistics corresponding to Supplementary Figure 5 on pRF size changes in eccentricity bin 4. P-values reflect whether t-test showed that subject average values were different from 0, for each ROI. Single, double and triple asterisks indicate uncorrected significance of <.05, <.01 and <.001 respectively. Right most columns indicate in how many out of 5 subjects the bootstrap test over voxels was different from 0 with uncorrect p-value of < .05.

**Table 22:** Statistics corresponding to Supplementary Figure 6 on correlations between eccentricity and size changes. P-values are uncorrected two-tailed tests whether t-test of over subject pRF eccentricity and size change correlations is different from 0. Single and triple asterisks indicate uncorrected significance of <.01 and <.001 respectively. Right most columns indicate in how many out of 5 subjects the bootstrap test over voxels was different from 0 with uncorrect p-value of < .05.

| ROI | R | N | t | p | pos in | neg in |
| --- | --- | --- | --- | --- | --- | --- |
| V1 | 0.461 | 5 | 2.888 | .045* | 2 | 0 |
| V2 | 0.704 | 5 | 6.411 | .003** | 4 | 0 |
| V3 | 0.704 | 5 | 7.219 | .002** | 5 | 0 |
| hV4 | 0.749 | 5 | 10.609 | <.001*** | 5 | 0 |
| VO | 0.886 | 5 | 12.673 | <.001*** | 5 | 0 |
| LO | 0.847 | 5 | 8.074 | .001** | 5 | 0 |
| V3AB | 0.861 | 5 | 6.954 | .002** | 5 | 0 |
| IPS0 | 0.500 | 5 | 2.450 | .070 | 3 | 0 |
| MT+ | 0.745 | 5 | 6.261 | .003** | 4 | 0 |
| combined | 0.799 | 5 | 5.431 | .006** | 4 | 0 |

**Table 23:** Statistics corresponding to Supplementary Figure 10. Astrisk indicate whether the average feature AMI over subjects was different from 0 with p < .05. Right most columns indicate in how many out of 5 subjects the bootstrap test over voxels was different from 0 with uncorrect p-value of < .05.

| ROI | N | mean diff | t | p | cohen d | pos in | neg in |
| --- | --- | --- | --- | --- | --- | --- | --- |
| V1 | 5 | 0.065 | 4.008 | .016* | 2.004 | 4 | 0 |
| V2 | 5 | 0.040 | 4.195 | .014* | 2.097 | 4 | 0 |
| V3 | 5 | 0.028 | 1.985 | .118 | 0.992 | 2 | 0 |
| hV4 | 5 | 0.055 | 3.235 | .032* | 1.618 | 4 | 0 |
| VO | 5 | 0.079 | 1.594 | .186 | 0.797 | 2 | 0 |
| LO | 5 | 0.026 | 1.122 | .325 | 0.561 | 2 | 1 |
| V3AB | 5 | 0.111 | 3.243 | .032* | 1.621 | 4 | 0 |
| IPS0 | 5 | 0.113 | 2.218 | .091 | 1.109 | 2 | 0 |
| MT+ | 5 | 0.060 | 1.531 | .201 | 0.766 | 3 | 0 |

**Table 24:**
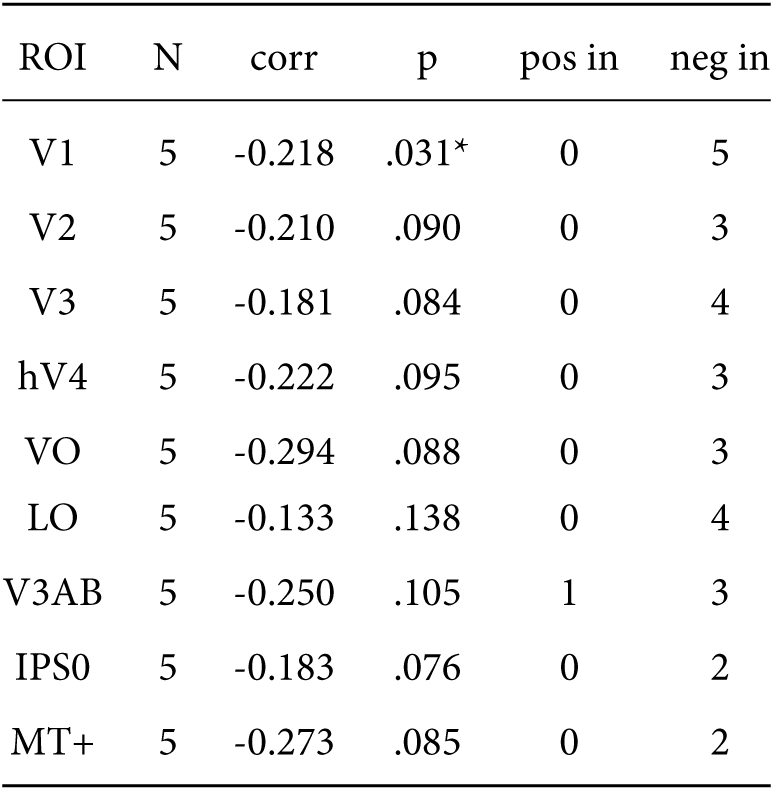
Statistics corresponding to Supplementary Figure 11. Astrisk indicate whether the spearman correlation between feature preference and eccentricity over subjects was different from 0 with p < .05. Right most columns indicate in how many out of 5 subjects the bootstrap test over voxels was different from 0 with uncorrect p-value of < .05.

